# Progressive deterioration of adaptive immune repertoires in Down syndrome linked to interferon hyperactivity and lymphoid tissue disorganization

**DOI:** 10.64898/2026.07.30.741516

**Authors:** Micah G. Donovan, Neetha Paul Eduthan, Brian F. Niemeyer, Elena Woods, Eric Hoffmeyer, Natalia Jaeger, Eleanor C. Britton, Bartosz B. Grzywacz, Joseph J. Fernandez, Galileo Dumont, Brian W. Herrmann, Norman R. Friedman, Dallas Jones, Zdenek Andrysik, Angela L. Rachubinski, Kelly D. Sullivan, Matthew D. Galbraith, Michael R. Verneris, Joaquin M. Espinosa

**Author notes:** Equal contributions, order alphabetically (Micah G. Donovan, Neetha Paul Eduthan, Brian F. Niemeyer, Elena Woods). Correspondence to | |.

## Abstract

Persons with Down syndrome (DS), the genetic condition caused by trisomy 21 (T21), display strong dysregulation of adaptive immunity, which underlies high risk of complications from infections, widespread autoimmunity, and poor vaccine responses. However, the mechanisms by which T21 dysregulates adaptive immunity across the lifespan remain poorly understood. We report here a multimodal analysis of adaptive immunity across development and aging in DS, including deep B cell profiling by mass cytometry matched to transcriptome and proteome data, B and T cell receptor sequencing (BCR, TCR), and spatial transcriptomics of tonsil tissue. T21 causes progressive shifts in B cell subsets together with accelerated age-dependent B cell loss linked to hyperactive interferon and JAK/STAT signaling. The peripheral immunoglobulin repertoire shows dysregulated class switching, progressive loss of diversity, differential VDJ usage, and imbalanced rates of somatic hypermutation across immunoglobulin isotypes mirrored by contraction and skewing of the TCR repertoire. In children with DS, tonsil tissues are highly disorganized with smaller germinal centers, fibrotic intrusions, lower rates of cell proliferation, strong inflammatory signaling, and transcriptional programs indicative of dysregulated lymphocyte homing and residence. Together, these results point to hyperactive interferon signaling as a driver of dysregulated adaptive immunity in DS amenable to early therapeutic intervention.

## INTRODUCTION

Down syndrome (DS), the medical condition caused by trisomy 21 (T21), involves widespread disruption of immune function across both innate and adaptive compartments^1,2^. A hallmark of this condition is sustained activation of interferon (IFN) signaling, stemming in part from increased dosage of IFN receptor genes encoded on chromosome 21 (*IFNAR1*, *IFNAR2*, *IFNGR2*, *IL10RB*), leading to an autoinflammatory state reminiscent of interferonopathies^3–5^. Within the innate immune system, heightened sensitivity to IFN stimulation associates with an inflammatory profile characterized by elevated production of many myeloid-derived cytokines and growth factors^6^. This underlying inflammatory tone associates with increased complications from infections, especially of the respiratory tract^7^, reflecting a possible combination of compromised immune defense and exacerbated inflammatory responses of pathogenic potential. In fact, severe complications from lung infections are a leading cause of morbidity and mortality in this population across the lifespan^7^. Within the adaptive immune system, persons with DS frequently show abnormalities in lymphocyte composition, including various imbalances in B and T subsets^1,2,8^, weaker vaccine responses^9^, and impaired thymic function^10^. Rates of autoimmune disorders, including autoimmune thyroid disease (AITD), celiac disease, type I diabetes, and various immune-mediated skin conditions, are significantly elevated^11,12^. Collectively, these findings highlight a dysregulated immune state marked by simultaneous immune overactivation and functional impairment with broad clinical consequences^13^.

B cell dysregulation is a consistent feature of DS, reflecting defects in development, subset distribution, and function^1,2^. Persons with DS commonly exhibit reduced total B cell numbers, with a marked decrease in naïve memory B cells, alongside an expansion of differentiated B cell subsets, age-associated B cells (ABCs), and plasmablasts^1,14,15^. Functionally, individuals with DS frequently exhibit impaired humoral vaccine responses, including lower antigen-specific antibody titers and accelerated waning of protective antibodies following immunization^9^. Conversely, there is also an increased propensity for autoantibody production^2,12^, consistent with the higher prevalence of autoimmune conditions. This combination of impaired protective humoral immunity and increased autoreactivity reflects a loss of B cell tolerance, likely influenced by chronic IFN signaling and dysregulated T cell crosstalk^1,2,8^.

T cell dysregulation is also obvious in DS. Individuals with DS commonly exhibit reduced total T cell numbers and decreased CD4+/CD8+ ratios^8^. CD4+ T cells display increased differentiation, polarization toward the Th1 and Th1/17 states, and overproduction of cytokines such as IL-17A, IL-22, IL-10, and MIP-3a^8^. CD8+ T cells are depleted of naïve subsets and enriched for effector, memory, and exhausted subsets, express higher levels of markers of activation and senescence (e.g., IFN-γ, Granzyme B, PD-1, KLRG1), and overproduce cytokines such as TNF-a, IFN-g, IL-2, MIP-1a, and IL-8^8^. Although frequencies of T regulatory cells (Tregs) are increased in DS, effector T cells are resistant to Treg-mediated suppression^8^. These changes are associated with hypersensitivity to TCR activation^8^, elevated baseline JAK/STAT signaling^1,8^, and hypersensitivity to IFN stimulation^1,8^.

Despite these advances, the mechanisms by which T21 disrupts adaptive immunity remain incompletely understood. For example, the temporal dynamics of B and T cell dysregulation across the lifespan in DS are unclear. Whereas some studies indicate fundamental impairments in B cell differentiation that may start *in utero*^16^, others indicate that B cell dysregulation is part of a broader immunosenescence phenotype associated with accelerated aging in DS^17^. It is also unclear the degree to which T21 dysregulates B and T cell function through cell-autonomous versus cell-extrinsic mechanisms. Given that B cell development and maturation is highly dependent on heterotypic interactions with antigen-presenting cells, T cells, and stromal cells in secondary lymphoid tissues, T21 could impact B cell development and function through multiple, non-mutually exclusive mechanisms. Likewise, dysregulation of the B cell lineage can alter T cell development, activation, and homeostasis through abnormalities in B cell-mediated antigen presentation, germinal center and T:B border interactions important for T follicular helper (Tfh) differentiation, and modulation of excessive T cell responses, among other possible mechanisms.

Within this context, we present here an integrated, orthogonal, multi-platform investigation of dysregulated adaptive immunity across the lifespan in DS with an emphasis on B cell dysfunction. Our approach combined high-dimensional mass cytometry of B cell populations with matched transcriptomic and proteomic analyses, alongside B cell receptor (BCR) and T cell receptor (TCR) sequencing, and spatial transcriptomic analysis of tonsillar tissue. Our lifespan analysis demonstrates a gradual redistribution of B cell subsets together with an accelerated age-associated decline in total B cell numbers in individuals with exacerbated IFN/JAK/STAT signaling. Analysis of circulating immunoglobulin sequences reveals a steady contraction of repertoire diversity, altered VDJ gene usage, and imbalanced rates of somatic hypermutation across isotype classes, mirrored by progressive impoverishment and skewing of the TCR repertoire. Consistent with these systemic findings, tonsillar architecture is profoundly disrupted in children with DS, as evidenced by reduced germinal center size, fibrotic architecture, diminished cellular proliferation, robust inflammatory signaling, and signatures of dysregulated lymphocyte migration and residence. Collectively, these results point to sustained IFN/JAK/STAT signaling as a driver of progressive dysregulation of adaptive immune repertoires in DS, with clear implications for therapeutic strategies aimed at restoring immune homeostasis from an early age.

## RESULTS

### Trisomy 21 causes progressive changes in the B cell lineage across the lifespan

Previous reports have documented dysregulation of the B cell lineage in DS^1,2^. However, the temporal effects of T21 on B cell function at different life stages are poorly understood. To investigate the effects of T21 on B cells across the lifespan, we completed a sliding-window analysis^18^ of mass cytometry data from 388 research participants (291 with DS versus 97 euploid controls) collected through the Human Trisome Project cohort study (HTP) (**Supplementary Fig. 1a-c**, see **Methods**). We employed 10-year windows progressively shifted by 1-year increments to identify temporal changes in B cell marker expression and subset frequencies (**Fig. 1a-b, Supplementary Fig. 1d, Supplementary Table 1**). Importantly, this mass cytometry dataset spans ages 1 to ∼60 years old, thus enabling a dissection of T21-associated effects from early childhood through late adult life (**Supplementary Fig. 1b**).

**Fig. 1.**
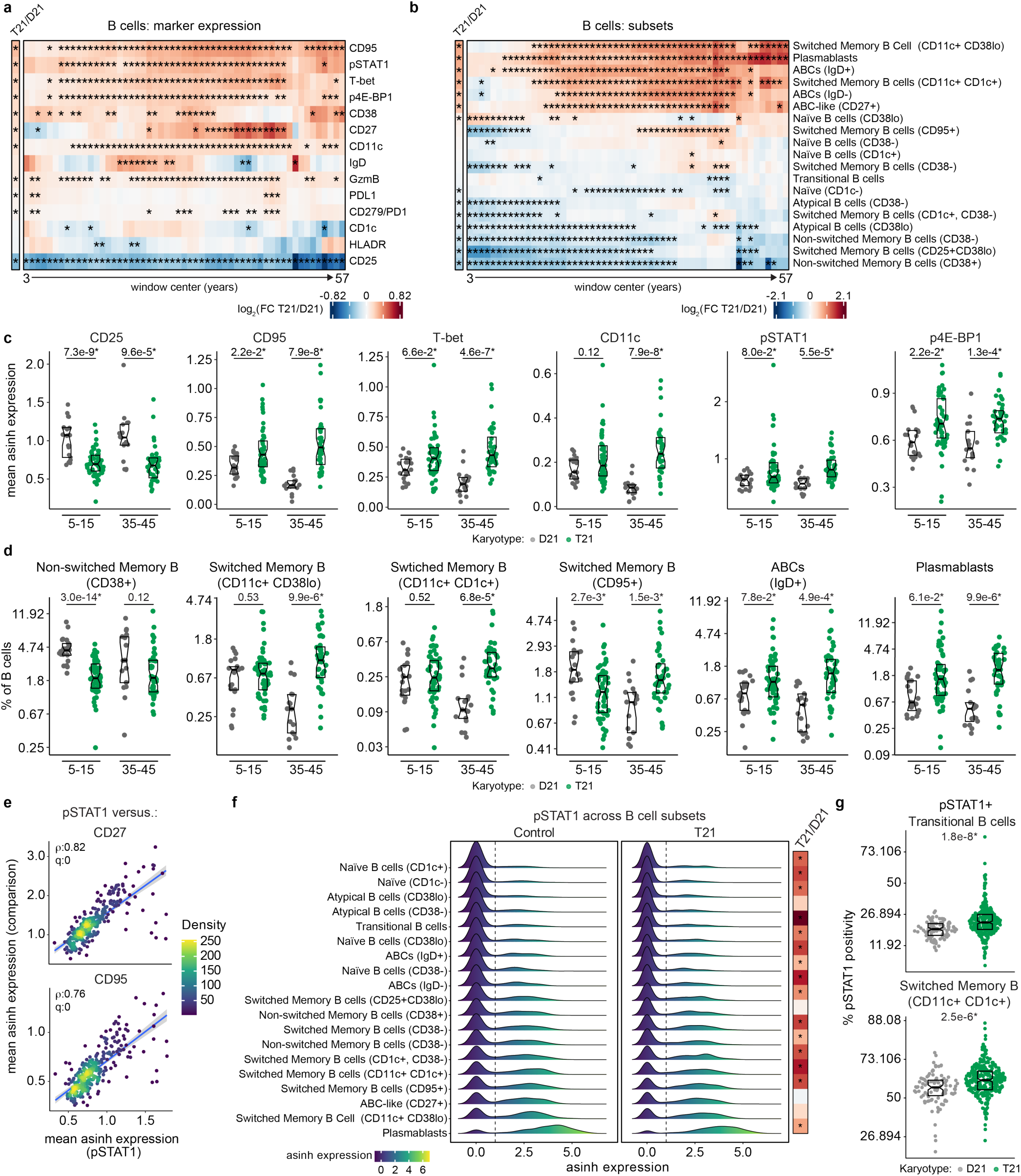
| Trisomy 21 causes progressive dysregulation of the B cell lineage across the lifespan. **a-b** Heatmaps showing results from differential sliding-window analysis (DESWAN) of (a) B cell marker expression and (b) B cell subsets from immune cell mass cytometry data (CyTOF) comparing individuals with trisomy 21 (T21, N total=292) to euploid controls (D21, N total=96). Comparisons were made within 10-year sliding-windows across the lifespan from 3 to 57 years of age. Color scales represent transformed (log2) fold changes comparing T21 to D21. Comparisons were made using linear regression (B cell markers) and beta regression (B cell subsets) applying Benjamini-Hochberg multiple hypothesis correction. Asterisks indicate significance (q<0.1). **c-d** Sina plots showing distributions of select (c) B cell markers and (d) B cell subsets across D21 and T21 in individuals aged 5-15 years (N D21/T21=21/53) and 35-45 years (N D21/T21=17/40). Statistics above data swarms indicate q-values from linear regression (B cell markers) and beta regression (B cell subsets) analyses applying the Benjamini-Hochberg method. **e** Scatter plots showing associations between expression of phosphorylated STAT1 (pSTAT1) vs. (top) CD27 and (bottom) CD95 expression among individuals with T21 (N=292). Points represent individual samples (colored by density) with lines indicating fitted regression trends with shaded 95% confidence intervals. Data are shown as mean inverse hyperbolic sine (asinh)-transformed expression. **f** Ridgeline density plots showing the distribution of asinh-transformed pSTAT1 expression across B cell subsets, stratified by D21 (N=96) and T21 (N=292). Vertical dashed lines indicate threshold for pSTAT1 positivity. Heatmap on the right displays results from beta regression comparing T21 vs. D21 for pSTAT1 positivity with color scales representing log2(fold change) and asterisks indicating significance (q<0.1). **g** Sina plots showing distributions of the percentage of pSTAT1 positive (top) transitional B cells and (bottom) CD11c+ CD1c+ switched memory B cells across D21 (N=96) and T21 (N=292). For sina plots (c-d, g) points represent individual samples (D21, gray; T21, green); boxes indicate medians and interquartile ranges, with notches approximating 95% confidence intervals.

**Supplementary Fig. 1.**
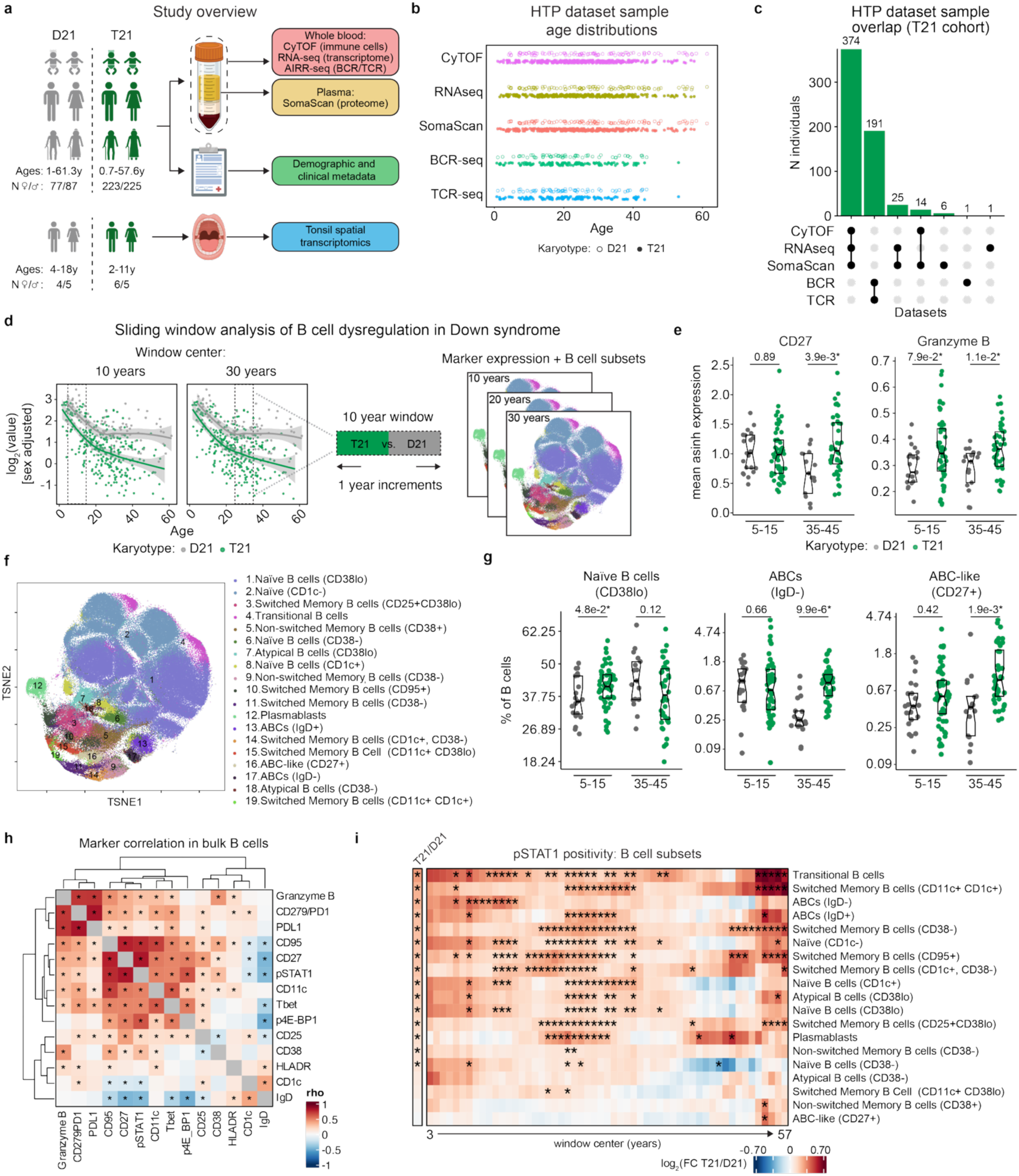
| Trisomy 21 causes progressive dysregulation of the B cell lineage across the lifespan. **a** Graphical summary of study design. **b** Age distribution of participants across datasets. Each point represents one individual. Open circles indicate euploid control (D21) participants and filled circles indicate individuals with trisomy 21 (T21). **c** UpSet plot displaying the overlap of multi-omic data generation from the same blood samples from individuals with T21. **d** Schematic of differential sliding-window analysis (DESWAN) approach to analyze differences in B cell marker expression and B cell subset frequencies between D21 and T21 across the lifespan. Scatter plots on the left show total B cell levels as a function of age in T21 (green) and D21 (gray) with vertical dashed lines representing boundaries of 10-year sliding windows. Solid lines represent localized polynomial (loess) fit lines, with 95% confidence intervals in gray. Within each sliding-window, B cell marker expression and B cell subset levels are compared between T21 and D21, shown by representative t-distributed stochastic neighbor embedding (t-SNE) plots. **e** Sina plots showing distributions of select B cell markers across D21 and T21 in individuals aged 5-15 years (N D21/T21=21/53) and 35-45 years (D21/T21=17/40). Statistics above data swarms indicate q-values from linear regression analysis applying the Benjamini-Hochberg multiple hypothesis correction. **f** tSNE plot showing immune cell mass cytometry (CyTOF) data projected into two-dimensional space (tSNE1 and tSNE2), with cells colored by annotated B cell subset identity. Each point represents an individual cell. **g** Sina plots showing distributions of select B cell subsets across D21 and T21 in individuals aged 5-15 years (N D21/T21=21/53) and 35-45 years (D21/T21=17/40). Statistics above data swarms indicate q-values from beta regression analysis applying the Benjamini-Hochberg multiple hypothesis correction**. h** Heatmap displaying Spearman correlations between B cell markers among bulk B cells across individuals with T21. Color scales indicate Spearman rho values with asterisks indicating significance (q<0.1) after Benjamini-Hochberg multiple hypothesis correction. **i** Heatmap showing results from beta regression analysis of phosphorylated STAT1 (pSTAT1, Tyr 701) positive B cell subsets comparing individuals with T21 vs. D21 in (left) the full cohort and (right) across individual 10-year sliding windows. Color scales represent transformed (log2) fold-changes from beta regression analyses with asterisks indicating significance (q<0.1) after Benjamini-Hochberg multiple hypothesis correction.

First, we analyzed various markers relevant to B cell function in the entire B cell population, which revealed temporally dynamic changes relative to age-matched euploid controls. B cells from young children with DS display decreased expression of the IL-2 receptor alpha chain (IL-2Rα/CD25), relative to children without DS (**Fig. 1a, c**). B cells expressing CD25 are poised to proliferate and acquire immunomodulatory features^19^. Lower expression of CD25 is consistently observed across the lifespan in DS, even in older adults, and may indicate an overall lower proliferative capacity (**Fig. 1a, c**).

As children with DS age, the peripheral B cell lineage displays progressively increased expression of CD95 (Fas receptor), T-bet (TBX21), and CD11c (ITGAX) relative to euploid controls (**Fig. 1a, c**). All three markers decrease with age in the control group, but increase with age in DS, leading to much exacerbated differences in adults (**Fig. 1c**). CD95 is expressed on activated B cells and mediates elimination of B cells that are autoreactive, low-affinity, or no longer needed after an immune response^20^. T-bet is a transcription factor upregulated in B cells by signals such as IFNγ and TLR activation^21^. CD11c is an integrin α chain that mediates cell adhesion and migration, facilitating antigen presentation and localization within inflamed tissues^21^. Together, elevation of these three markers points to sustained activation and differentiation, heightened susceptibility to activation-induced cell death, and a tissue-homing phenotype associated with chronic immune stimulation. Notably, steady increases in these markers correlate with elevated phosphorylation of STAT1 (pSTAT1) and 4E-BP1 (p4E-BP1), indicative of increased JAK/STAT and mTOR signaling, respectively (**Fig. 1a, c**). In B cells, pSTAT1 promotes enhances antigen presentation and supports differentiation toward specialized subsets^22^. The mTORC1/4E-BP1/eIF4E axis promotes antibody class switching^23^ and elevated p4E-BP1 is associated with differentiation into antibody-secreting plasmablasts and plasma cells, which require high rates of protein synthesis^23^.

Later on, well into adult life, B cells from persons with DS display increased expression of CD27 (**Fig. 1a**, **Supplementary Fig. 1e**), a member of the TNFR superfamily and a key marker of antigen-experienced B cells^24^. CD27 is most prominently expressed on memory B cells and subsets of plasmablasts, distinguishing them from naïve B cells. CD27⁺ B cells are capable of rapid and robust antibody production upon antigen re-exposure, making them critical for long-term humoral immunity. Notably, we also observed higher expression of Granzyme B on B cells from people with DS, which has been associated with both cytotoxic B cells^25^ and control of T cells^26^ (**Fig. 1a**, **Supplementary Fig. 1e**).

Next, we completed sliding-window analysis of mass cytometry data to identify differences in subset frequencies across the lifespan in DS. Toward this end, we performed unsupervised clustering and visualization, followed by expert curation and naming, to identify 19 distinct B cell subsets (**Supplementary Fig. 1f**, **Supplementary Table 1**, see **Methods**). Beta regression testing of differential cell frequencies revealed clear life stage-specific events (**Fig. 1b, d**). Young children with DS display significantly lower frequencies of non-switched memory B cells subsets (both CD38+ and CD38-) (**Fig. 1b, d**). Because non-switched memory B cells, which have encountered antigen but have not undergone class-switch recombination^27^, can re-enter germinal centers and undergo further maturation^28^, their depletion in young children with DS could impair early humoral responses and contribute to increased susceptibility to infections.

Interestingly, as children with DS age, they display a progressive increase in the frequencies of switched memory B cells expressing CD11c (CD11c+ CD38lo, CD11c+ CD1c+), and, later in adult life, subsets expressing CD95 relative to euploid controls (**Fig. 1b, d**). These subsets decrease in frequency with age in the euploid cohort, but not in the T21 cohort, leading to their over-representation in DS. This profile has been observed under IFN-γ–dominated conditions, including chronic infections (e.g., malaria, HIV), autoimmune diseases (e.g., systemic lupus erythematosus – SLE), and inflammaging^29,30^.

Starting in adolescence and progressing into adult life, the B cell lineage of persons with DS is marked by progressively higher frequencies of ABC subsets and plasmablasts relative to their age-matched euploid controls (**Fig. 1b, d, Supplementary Fig. 1g**). These observations align with earlier reports demonstrating skewing toward antibody-producing and autoreactive cells^1,2^, and further suggest that this process is progressive, with greater perturbation observed in adults with DS.

Given the prominent role of IFN/JAK/STAT signaling in B cell lineage development, we next investigated the interplay between elevated pSTAT1 and B cell dysregulation in DS. Correlation analysis shows that pSTAT1 levels are strongly associated with CD27, CD95, CD11c, and T-bet (**Fig. 1e, Supplementary Fig. 1h**). Across B cell subsets, pSTAT1 levels increase progressively, with the lowest signaling in naïve cells and a gradual elevation in switched memory B cells, ABCs, and plasmablasts, which exhibit the highest levels (**Fig. 1f**). Consistently, the fraction of pSTAT1 positive cells is significantly higher in DS across most subsets (**Fig. 1f-g**). Sliding-window analysis of pSTAT1 positivity indicates that JAK/STAT signaling starts early in life and is maintained across the lifespan in DS (**Supplementary Fig. 1i**).

Altogether, these results demonstrate a progressive dysregulation of the B cell lineage associated with lifelong elevation of baseline JAK/STAT signaling in DS.

### Extreme B cell lymphopenia associates with elevated IFN/JAK/STAT signaling in Down syndrome

Although lower B cell counts are observed in persons with DS at all life stages, this phenomenon is exacerbated with age, albeit with strong inter-individual variability (**Fig. 2a**). At every life stage, some individuals with DS display extreme B cell lymphopenia, whereas others show B cell counts similar to age-matched euploid controls (**Fig. 2a**). To investigate sources of this heterogeneity across the lifespan, we calculated residuals from a loess fit generated for the T21 cohort and defined the top and bottom quartiles to create two groups of persons with DS: B^High^, who display frequencies of B cells similar to their age-matched controls, and B^Low^, who display extreme B cell depletion (**Fig. 2a-b**). We then performed a multi-omics comparison of the two groups, including analysis of their immune profiles, whole blood transcriptome, and plasma proteome.

**Fig. 2.**
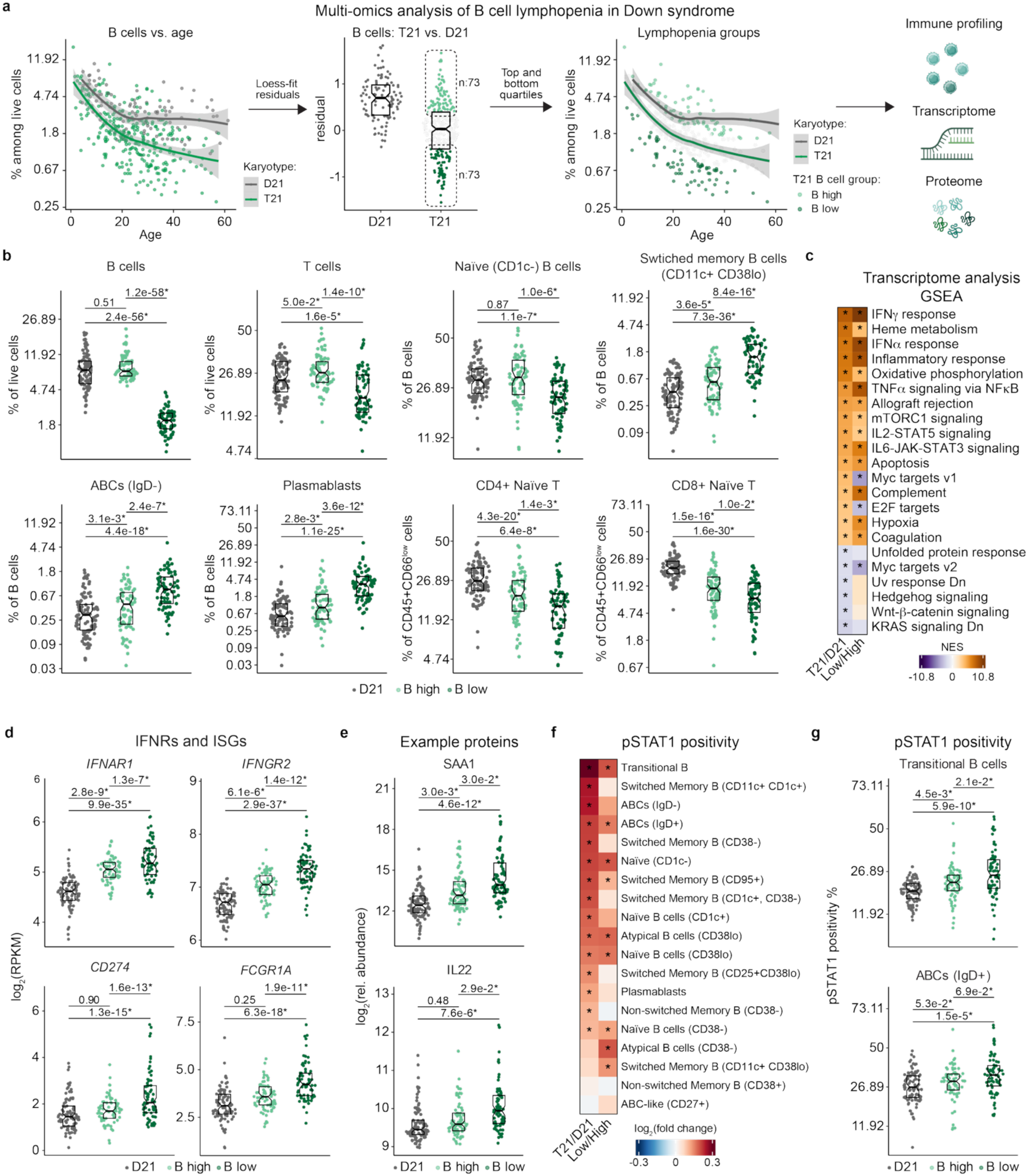
| B cell lymphopenia associates with hyperactive IFN/JAK/STAT signaling in Down syndrome. **a** Schematic outlining the multi-omic analysis framework used to investigate B cell lymphopenia in Down syndrome (DS). Localized polynomial regression (LOESS) was used to model total B cell abundance, measured by immune cell mass cytometry, as a function of age. Individuals with trisomy 21 (T21) were stratified based on residuals from the LOESS fit, with B cell low and B cell high groups defined as residuals below the first quartile (Q1) and above the third quartile (Q3), respectively. Multi-omic differences between B cell low and B cell high groups were evaluated across immune cell profiles, whole-blood transcriptome (RNA-seq), and plasma proteome. **b** Sina plots displaying the distribution of immune cell proportions among total live cells, total B cells, or total non-granulocytes (CD45+ CD66low) in euploid controls (D21; N=96) and individuals with T21 with high (B high; N=73) and low (B low; N=73) B cells as defined in (a). Differences between groups were determined by beta regression with Benjamini-Hochberg multiple hypothesis correction. Statistics above data swarms indicate q-values. **c** Heatmap displaying results from Gene Set Enrichment Analysis (GSEA) of whole-blood transcriptome comparing T21 (N=304) vs. D21 (N=90) and among individuals with T21 with low (N=71) vs. high (N=71) B cells. Color scale represents normalized enrichment score (NES), with asterisks indicating significance (q<0.1) after Benjamini-Hochberg multiple hypothesis correction. **d** Sina plots showing distributions of select interferon receptor (IFNR) gene and interferon-stimulated gene (ISG) expression from whole blood RNA-seq in D21 (N=90) and individuals with T21 with high (B high; N=71) and low (B low; N=71) levels of total B cells. Differences between groups were determined by DESeq2 with Benjamini-Hochberg multiple hypothesis correction. Statistics above data swarms indicate q-values. **e** Sina plots showing distribution of levels of select plasma proteins, measured by SomaScan, in D21 (N=96) and individuals with T21 with high (B high; N=73) and low (B low; N=73) levels of total B cells. Differences between groups were determined by linear regression with Benjamini-Hochberg multiple hypothesis correction. Statistics above data swarms indicate q-values. **f** Heatmap summarizing percentages of pSTAT1+ cells across the indicated B cell subsets. The two columns indicate comparisons of individuals with T21 (N=292) versus euploid controls (D21, n=96) (left) and individuals with T21 in the B cell low versus B cell high groups (N=73 per group). **g** Sina plot showing percentages of pSTAT1+ positive transitional B cells (top) and ABCs (IgD+) in euploid controls (D21, N=96) and individuals with T21 in the B cell low versus B cell high groups (N=73 per group). For sina plots (b, d, e, g), points represent individual samples (D21, gray; T21 B high, light green; T21 B low, dark green); boxes indicate medians and interquartile ranges, with notches approximating 95% confidence intervals.

**Supplementary Fig. 2.**
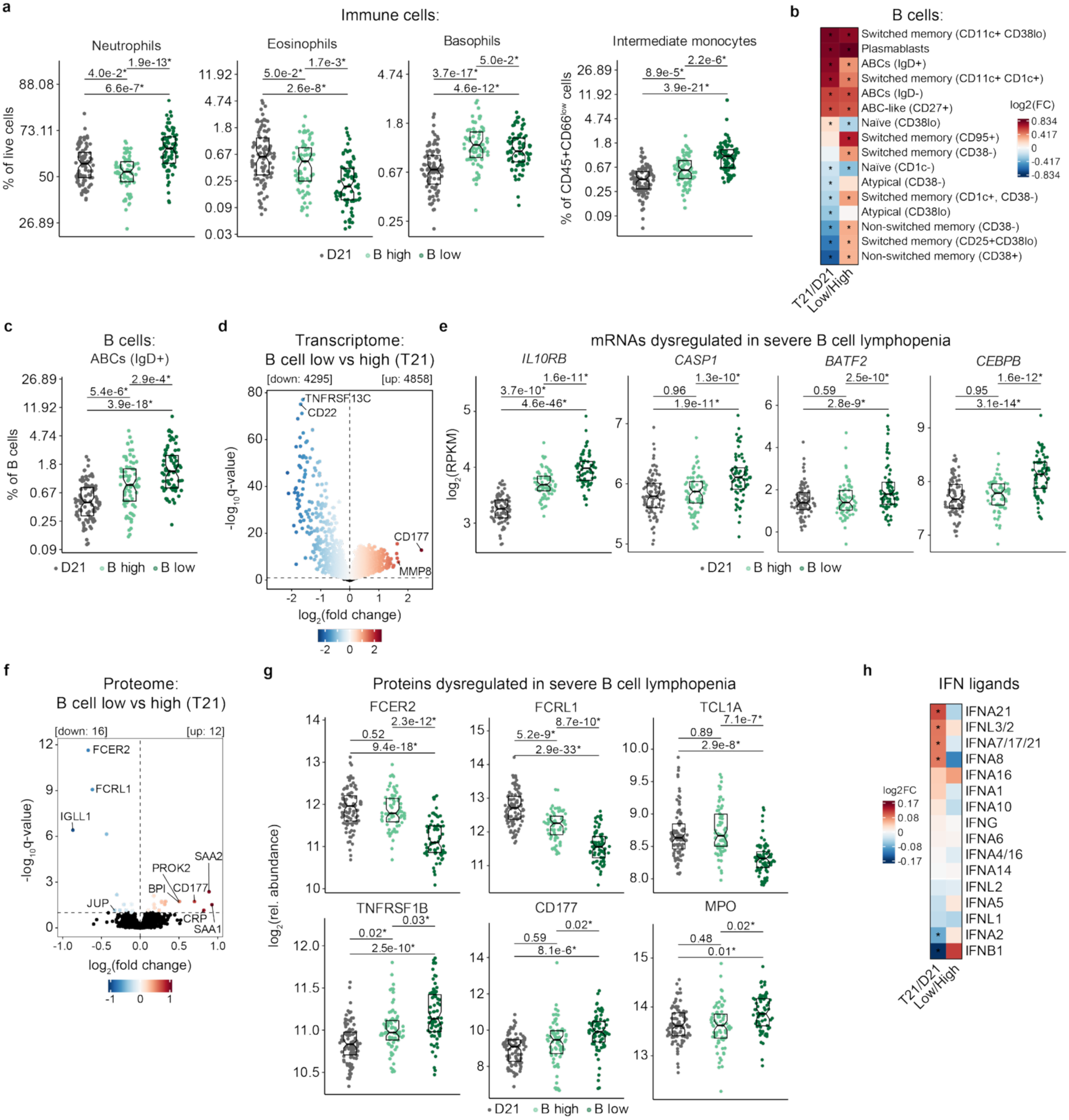
| Multi-omics analysis of B cell lymphopenia in Down syndrome. **a** Sina plots displaying the distribution of immune cell proportions among total live cells or non-granulocytes (CD45+ CD66low) in D21 (N=96) and individuals with T21 with high (B high; N=73) and low (B low; N=73) levels of total B cells. Differences between groups were determined by beta regression with Benjamini-Hochberg multiple hypothesis correction. Statistics above data swarms indicate q-values. **b** Heatmap displaying differences among B cell subsets comparing T21 (N=292) vs. D21 (N=96) and individuals with T21 with low (N=73) vs. high (N=73) B cells. Differences between groups were determined by beta regression with Benjamini-Hochberg multiple hypothesis correction. Color scales indicate transformed (log2) fold-changes between groups. Asterisks indicate significance (q<0.1). **c** Sina plots displaying the distribution of ABC (IgD+) subset proportions in D21 (N=96) and individuals with T21 with high (B high; N=73) and low (B low; N=73) levels of total B cells. Differences between groups were determined by beta regression with Benjamini-Hochberg multiple hypothesis correction. Statistics above data swarms indicate q-values. **d** Volcano plot of whole blood RNA differential gene expression comparing individuals with T21 with low (N=71) vs. high (N=71) levels of total B cells. Differential expression was analyzed by DESeq2 with Benjamini-Hochberg multiple hypothesis correction. Points are colored by log2(fold-change) with black points representing non-significant differences (q≥0.1). **e** Sina plots showing distributions of select interferon-stimulated gene (ISG) expression from whole blood RNA-seq in D21 (N=96) and individuals with T21 with high (B high; N=71) and low (B low; N=71) levels of total B cells. Differences between groups were determined by DESeq2 with Benjamini-Hochberg multiple hypothesis correction. Statistics above data swarms indicate q-values. **f** Volcano plot of plasma differential protein abundance comparing individuals with T21 with low (N=73) vs. high (N=73) levels of total B cells. Differential abundance was analyzed by linear regressions with Benjamini-Hochberg multiple hypothesis correction. Points are colored by log2(fold-change) with black points representing non-significant differences (q≥0.1). **g** Sina plots showing distribution of levels of select plasma proteins in D21 (N=96) and individuals with T21 with high (B high; N=73) and low (B low; N=73) levels of total B cells. Differences between groups were determined by linear regression with Benjamini-Hochberg multiple hypothesis correction. Statistics above data swarms indicate q-values. **h** Heatmap displaying differential abundance of interferon (IFN) ligands comparing D21 versus T21 and individuals with T21 with low versus high levels of total B cells. Color scale indicates log2(fold-change) from linear regression analyses with Benjamini-Hochberg multiple hypothesis correction. Asterisks indicate significance (q<0.1). For sina plots (a, c, e, g), points represent individual samples (D21, gray; T21 B high, light green; T21 B low, dark green); boxes indicate medians and interquartile ranges, with notches approximating 95% confidence intervals.

Among major immune cell lineages, the B^Low^ group displays lower T cell counts, elevated neutrophils, and lower eosinophils and basophils (**Fig. 2b, Supplementary Fig. 2a**). Among B cell subsets, those in the B^Low^ group show many significant differences, including fewer naïve B cell subsets, increased frequencies of multiple memory subsets (both non-switched and switched), and supra-elevated levels of ABC subsets and plasmablasts (**Fig. 2b, Supplementary Fig. 2b-c**). Thus, B cell lymphopenia in DS is clearly associated with strong shifts away from naïve states toward antigen-experienced and antibody-producing cells. The B^Low^ group also shows depletion of naïve CD4+ and CD8+ subsets, as well as elevated intermediate monocytes (**Fig. 2b**, **Supplementary Fig. 2a**). Clearly, B cell depletion is accompanied by global immune remodeling in DS.

Analysis of the whole blood transcriptome revealed vast gene expression changes in the B^Low^ group (**Supplementary Fig. 2d**). Gene Set Enrichment Analysis (GSEA) revealed that extreme lymphopenia associates with exacerbated IFNγ and IFNα responses, along with enrichment of accompanying pro-inflammatory programs (e.g., Inflammatory Response, TNFα signaling, Allograft Rejection signature, mTORC1 signaling, IL-2 STAT5 signaling, IL-6 JAK STAT3 signaling) and depletion of hallmark signatures associated with cell proliferation (Myc Targets V1 and V2, E2F targets) (**Fig. 2c**). Hyperactive IFN signaling in the B^low^ group associates with elevated expression of the four IFNRs encoded on chr21 as well as dozens of ISGs, including many with known roles in control of B cell function, such as *CD274*, *FCGR1A*, *CASP1*, *BATF2*, and *CEBPB* (**Fig. 2d**, **Supplementary Fig. 2e**).

Analysis of the plasma proteome revealed relatively fewer changes. The B^low^ group shows strong depletion of FCER2 (the low affinity receptor for IgE), FCRL1 (the Fc Receptor Like 1), IGLL1 (Immunoglobulin Lambda Like Polypeptide 1, the light chain of the preB cell receptor), and TCL1A (T Cell Leukemia/Lymphoma 1A, a coactivator of the cell survival kinase AKT in T cells) (**Supplementary Fig. 2f-g**). Conversely, the B^low^ group shows marked elevation of multiple markers of systemic inflammation (CRP, SAA1, SAA2), elevated IL-22, higher levels of soluble the TNFR2 subunit TNFRSF1B, and higher levels of multiple neutrophil-derived proteins, such as CD177, BPI (Bactericidal Permeability Increasing Protein), and MPO (myeloperoxidase) (**Fig. 2e**, **Supplementary Fig. 2f-g**). Notably, extreme B cell lymphopenia is not associated with elevated levels of the IFN ligands, confirming the notion that elevated IFN signaling is driven mostly by IFNR overexpression in DS^3,5^, not by IFN ligand over-production (**Supplementary Fig. 2h**). Analysis of pSTAT1 positivity across B cell subsets revealed consistently elevated signals in the B^low^ group, particularly among naïve, transitional, atypical, switched memory, and ABC subsets (**Fig. 2f-g**).

Taken together, these results point to hyperactive IFN/JAK/STAT signaling as a potential driver of B cell lymphopenia in DS.

### The B cell receptor repertoire is progressively restricted and skewed in Down syndrome

Next, we analyzed the BCR repertoire in 192 research participants, 132 with DS versus 60 age– and sex-matched euploid controls through sequence analysis of BCR heavy chains using whole blood RNA samples (see **Methods**). First, we examined isotype frequencies, which revealed significantly lower levels of IgM and IgD sequences, along with significantly elevated usage of IgG and IgA (**Fig. 3a**). This phenomenon results in highly elevated IgG/IgM and IgG/IgD ratios in DS (**Fig. 3b**). Intriguingly, we also observed lower usage of IgE sequences (**Fig. 3a**), which could be explained by the known roles of Type II IFN signaling on suppression of IgE class switching^31^. In fact, IgE is one of the most depleted proteins in the circulating proteome of people with DS^6^, which could explain their protection from allergic sensitization^32^.

**Fig. 3.**
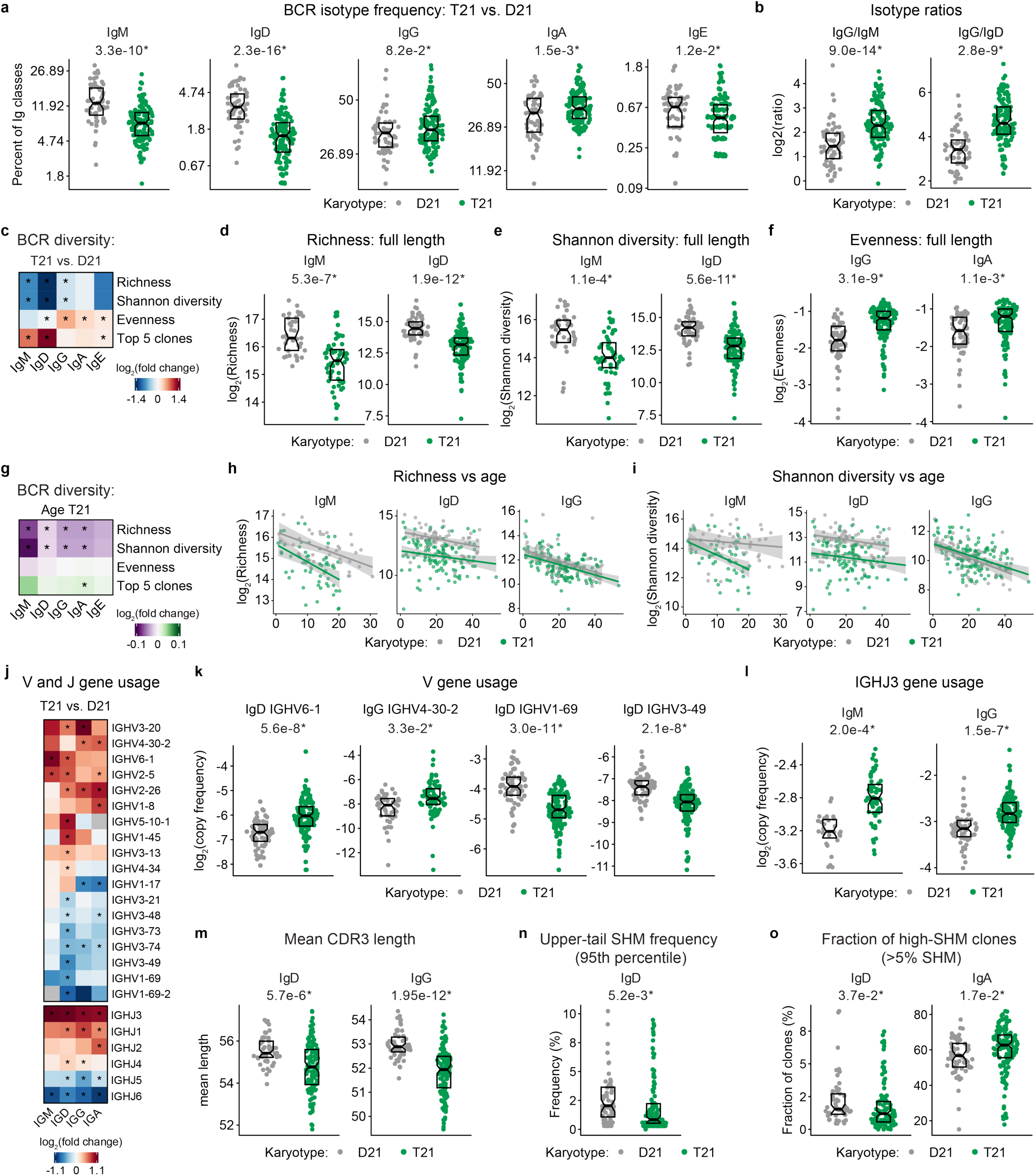
| The B cell receptor repertoire is highly restricted and skewed in Down syndrome. **a** Sina plots showing Ig class isotype frequency across euploid controls (D21, N=60) and participants with trisomy 21 (T21, N=132). **b** Sina plots showing ratios of Ig class isotype frequency across D21 (N=60) and participants with T21 (N=132). **c** Heatmap showing differences in B cell receptor sequencing (BCR-seq) diversity metrics across Ig isotypes comparing individuals with T21 (N=132) vs. D21 (N=60). **d-f** Sina plots showing for select Ig class isotypes (d) clone richness, (e) Shannon diversity, and (f) evenness comparing individuals with T21 vs. D21. Diversity metrics were calculated per Ig class on samples with at least 30 clones using Hill diversity numbers, where the parameter q determines sensitivity to clone frequency. D21/T21 samples sizes were: IgM (37/59), IgD (58/123), IgG (60/132), IgA (60/132). **g** Heatmap showing the effect of age on B cell repertoire diversity metrics across Ig class isotypes in individuals with T21. Sample sizes as in d-f. **h-i** Scatter plots showing B cell repertoire diversity metrics for select Ig class isotypes as a function of age in individuals with T21 vs. D21. Sample sizes as in d-f. **j** Heatmap showing differences in V and J gene usage across Ig class isotypes comparing individuals with T21 vs. D21. Sample sizes as in d-f. **k-l** Sina plots showing the distribution of select V (k) and J (l) gene copy frequency across select Ig class isotypes in D21 and T21. Sample sizes as in d-f. **m** Sina plots showing the distribution of mean Complementarity-Determining Region 3 (CDR3) tail lengths across IgM and IgD isotypes in D21 (N=60) and T21 (N=132). **n-o** Sina plots comparing somatic hypermutation (SHM) across D21 (N = 60) and T21 (N = 132). (n) 95th percentile SHM frequency, representing the most highly mutated clones within each subject’s B cell repertoire. (o) Fraction of highly mutated clones (>5% SHM) within each subject’s B cell repertoire. Differences determined by two-sided Wilcoxon rank-sum test (Mann–Whitney U test). Panels a-o show BCR-seq of full-length fragments. Differences in isotype frequency (a) and top 5 clone frequency (c, g) were determined by beta regression with Benjamini–Hochberg multiple-testing correction. Differences in diversity metrics (c, d-i), V/J gene usage (j-l) and mean CDR3 tail length were determined by linear regression with Benjamini–Hochberg multiple-testing correction. For sina plots (a-b, d-f, k-o) points represent individual samples (D21, gray; T21, green); boxes indicate medians and interquartile ranges, with notches approximating 95% confidence intervals. For scatter plots (h-i) points represent individual samples with lines indicating fitted regression trends with shaded 95% confidence intervals. For heatmaps (c,g,j), color scales indicate log2(fold-change) with asterisks denoting significance (q<0.1).

**Supplementary Fig. 3.**
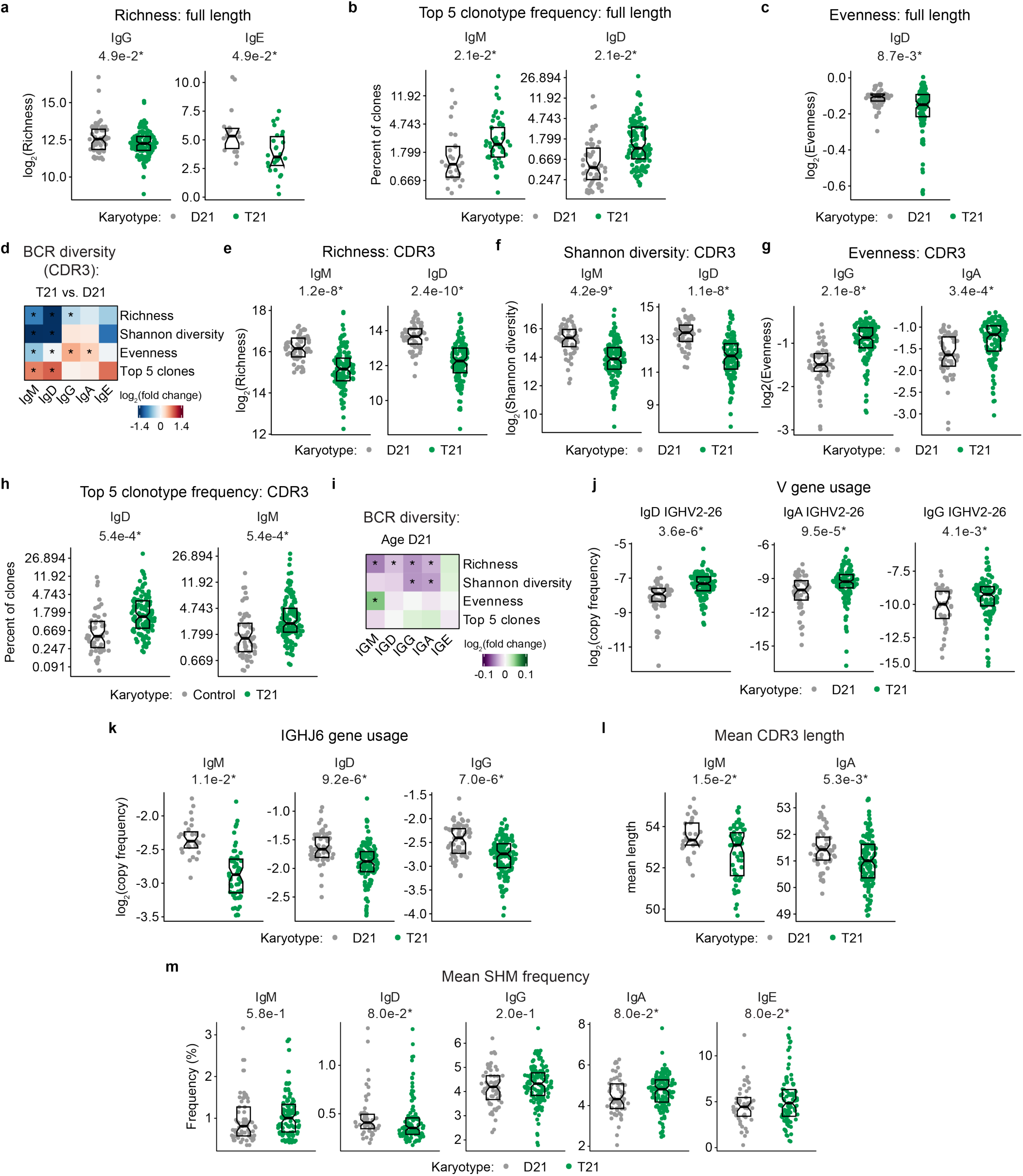
| The B cell receptor repertoire is highly dysregulated and skewed in Down syndrome. **a** Sina plots showing for select Ig class isotypes clone richness from BCR-seq of full-length fragments comparing individuals with trisomy 21 (T21; N IgG=132; N IgE=27) vs. euploid controls (D21; N IgG=60; N IgE=18). Diversity metrics were calculated per Ig class on samples with at least 30 clones using Hill diversity numbers. **b** Sina plots showing the cumulative frequency of top 5 clones for IgM and IgD isotypes across D21 (N IgM=37; N IgD=59) and T21 (N IgM=59; N IgD=125) from BCR-seq of full-length fragments. **c** Sina plot showing for IgD clone repertoire evenness from BCR-seq of full-length fragments across D21 (N=58) and T21 (N=123). **d** Heatmap showing differences in B cell receptor sequencing (BCR-seq) of Complementarity-Determining Region 3 (CDR3) regions across Ig isotypes comparing individuals with T21 (N=132) vs. D21 (N=60). **e-h** Sina plots showing for select Ig class isotypes (e) clone richness, (f) Shannon diversity, (g) evenness, and (h) top 5 clone frequency from BCR-seq of CDR3 regions comparing individuals with T21 vs. D21. Diversity metrics were calculated per Ig class on samples with at least 30 clones using Hill diversity numbers. D21/T21 samples sizes were: IgM (37/59), IgD (58/123), IgG (60/132), IgA (60/132). **i** Heatmap showing the effect of age on B cell repertoire diversity metrics across Ig class isotypes in euploid controls (D21, N=60). **j** Sina plots showing the distribution of select V gene copy frequency across select Ig class isotypes in D21 (N=60) and T21 (N=132) from BCR-seq of full-length fragments. **k** Sina plots showing the distribution of IGHJ6 copy frequency across select Ig class isotypes in D21 (N=60) and T21 (N=132). **l** Sina plots showing the distribution of mean Complementarity-Determining Region 3 (CDR3) tail lengths across IgM and IgD isotypes in D21 (N=60) and T21 (N=132). **m** Sina plots showing the distribution of mean somatic hypermutation (SHM) frequency across Ig isotypes in D21 (N=60) and T21 (N=132). Panels a-c and i-m show BCR-seq of full-length fragments. Panels d-h show BCR-seq of CDR3 regions. Differences in top 5 clone frequency (b,d,h-i) were determined by beta regression with Benjamini–Hochberg multiple-testing correction. Differences in diversity metrics (a,c,d-g, i) and V/J gene usage (j-k) were determined by linear regression with Benjamini–Hochberg multiple-testing correction. For sina plots (a-c,e-h,j-m), points represent individual samples (D21, gray; T21, green); boxes indicate medians and interquartile ranges, with notches approximating 95% confidence intervals.

We then analyzed BCR repertoire richness (number of unique clonotypes), evenness (uniformity in clonal distribution), Shannon diversity (a combined measure of richness plus evenness), and top 5 clonotype usage for each isotype separately, both for the full-length sequence and the Complementarity-Determining Region 3 (CDR3) sequence. This exercise revealed decreased richness and diversity along with increased top clonotype usage for IgM and IgD, concurrent with increased evenness for the IgG, IgA, and IgE isotypes (**Fig. 3c-f**, **Supplementary Fig. 3a-h**). Altogether, these results indicate immune skewing in DS marked by contraction of naïve subsets and expansion of class-switched, antigen-experienced populations, reminiscent of what has been observed in the context of chronic infections^33^, autoimmune conditions^34^, and conditions of chronic mucosal inflammation^35^. Notably, richness and diversity decrease with age for multiple isotype classes in both euploid controls and DS (**Fig. 3g-I, Supplementary Fig. 3i**).

Next, we investigated potential differences in IgH V and J gene usage, which revealed many sequences that are significantly over-represented or depleted in DS across IgM, IgD, IgG, and IgA isotypes (**Fig. 3j-l**, **Supplementary Fig. 3j-k**). Prominent examples of enriched V sequences in DS include *IGHV3-20* (previously associated with autoimmune aplastic anemia^36^), *IGHV4-30-2* (previously associated with rejection in kidney transplant patients^37^), *IGHV6-1* (previously found enriched in patients with lymphatic filariasis^38^), *IGHV2-5*, *IGHV1-8*, *IGHV5-10-1*, and *IGHV4-34* (previously found enriched in SLE^39^). Prominent examples of depleted sequences include *IGHV1-69*, and multiple *IGHV3* variants (–49, –74, –73, –48, –21), many of which were found decreased in SLE and other autoimmune conditions^40^. Analysis of J gene usage revealed consistently increased usage of J1-J4 sequences (shorter J segments), most prominently J3, concurrent with significant depletion of J5 and J6 sequences (longer J segments) (**Fig. 3j, l**, **Supplementary Fig. 3k**). This phenomenon demonstrates a shift toward shorter CDR3 regions in DS across multiple isotype classes (**Fig. 3m, Supplementary Fig. 3l**), a feature observed in many autoimmune conditions^41^.

We next assessed somatic hypermutation (SHM) across BCR isotypes. Mean SHM was significantly lower in IgD and higher in IgA and IgE in T21, with no significant differences observed for IgG or IgM (**Supplementary Fig. 3m**). Analysis of the upper tail of the SHM distribution (95th percentile SHM frequency) demonstrated reduced SHM among the most highly mutated IgD clones (**Fig. 3n**). Likewise, analysis of the fraction of highly mutated clones (>5% SHM) revealed fewer highly mutated IgD clones but a greater proportion of highly mutated IgA clones in T21 (**Fig. 3o**). These findings suggest isotype-specific alterations in affinity maturation, potentially reflecting reduced germinal center participation of IgD⁺ B cells alongside enhanced maturation of mucosal IgA responses.

Altogether, these results highlight the strong effects of T21 on B cell maturation and training, leading to significant dysregulation of humoral immunity in DS.

### Parallel dysregulation and skewing of the T cell receptor repertoire in Down syndrome

Given that B cell responses are tightly regulated by T cell help, particularly within germinal centers, changes in BCR repertoire architecture may arise from or be accompanied by abnormalities in the T cell compartment. Previous reports have documented extensive dysregulation of the T cell lineage in DS^1,2,8^, but differences in the TCR repertoire are less characterized. Therefore, we analyzed the TCR repertoire through sequencing of TCR alpha and beta chain genes (TRA, TRB). Both TRA and TRB display decreased richness, diversity, and evenness, along with increased representation of the top 5 clones in DS, with accelerated age-dependent deterioration of the TCR repertoire relative to euploid controls (**Fig. 4a-i**).

**Fig. 4.**
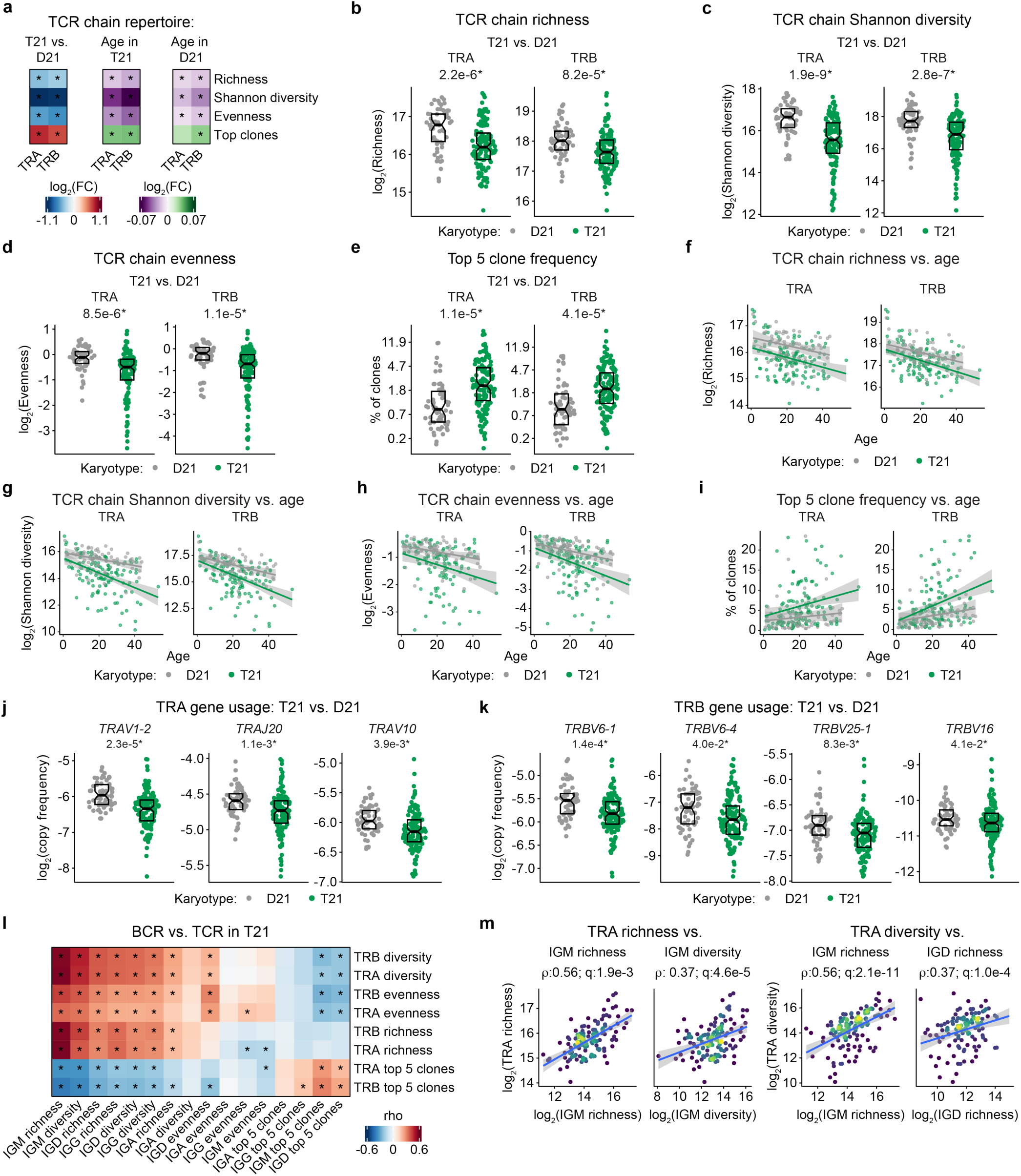
| Parallel dysregulation and skewing of the T cell receptor repertoire in Down syndrome. **a** Heatmap showing differences in T cell receptor sequencing (TCR-seq) diversity metrics across T cell receptor chains comparing individuals with T21 vs. D21 and age effects in both T21 and D21. **b-e** Sina plots showing for TCR chains (b) clone richness, (c) Shannon diversity, (d) evenness, and (e) top 5 clone frequency comparing individuals with T21 vs. D21. Diversity metrics were calculated per TCR chain on samples with at least 30 clones using Hill diversity numbers, where the parameter q determines sensitivity to clone frequency. **f-i** Scatter plots showing TCR repertoire diversity metrics as a function of age in individuals with T21 and D21. Points represent individual samples with lines indicating fitted linear regression trends with shaded 95% confidence intervals. **j-k** Sina plots showing the distribution of select V and J gene copy frequency across TCR alpha and beta chains in D21 and T21. **l** Heatmap displaying Spearman correlations between BCR and TCR diversity metrics in individuals with T21. Asterisks indicate significance (q-value < 0.1) after Benjamini-Hochberg multiple hypothesis correction. **m** Scatter plots showing associations between select B and T cell receptor diversity metrics in individuals with T21. Points represent individual samples with lines indicating fitted linear regression trends with shaded 95% confidence intervals. Panels a-m show TCR-seq of CDR3 regions in T21 (n=131) and D21 (n=60). Differences in diversity metrics (a-d) and V/J gene usage (j-k) were determined by linear regression with Benjamini–Hochberg multiple-testing correction. Differences in top 5 clone frequency (a,e) were determined by beta regression with Benjamini–Hochberg multiple-testing correction. For sina plots (b-e, j-k) points represent individual samples (D21, gray; T21, green); boxes indicate medians and interquartile ranges, with notches approximating 95% confidence intervals.

**Supplementary Fig. 4.**
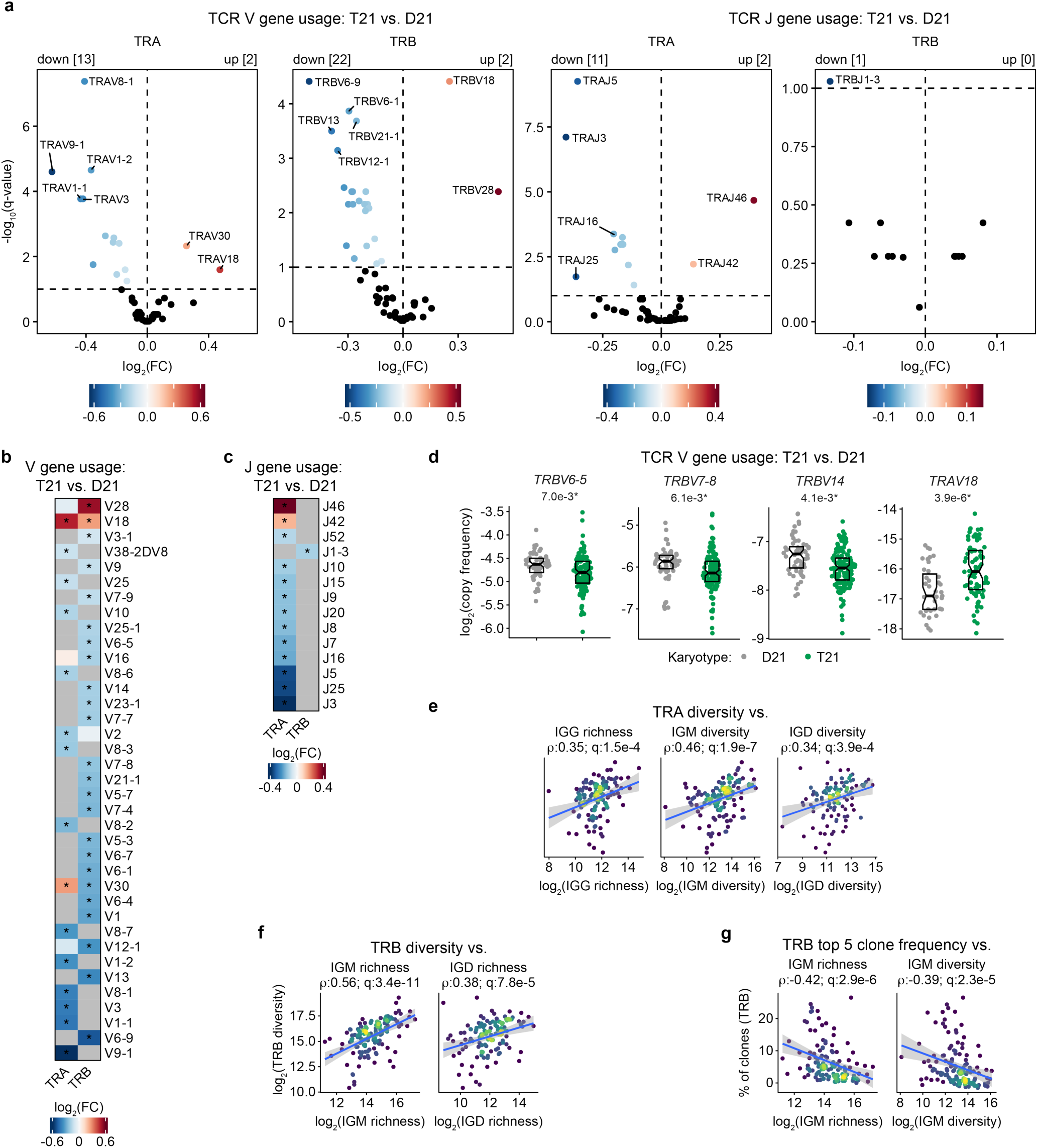
| Mirrored dysregulation and skewing of the T cell receptor repertoire in Down syndrome. **a** Volcano plots of differential V and J gene usage across T cell receptor (TCR) chains comparing individuals with T21 vs. D21. Differences determined by linear regression with Benjamini-Hochberg multiple hypothesis correction. Points are colored by log2(fold-change) with black points representing non-significant differences (q>0.1). **b-c** Heatmaps displaying differential V (b) and J (c) gene usage across alpha (TRA) and beta (TRB) chains. Examples selected from (a). Asterisks indicate significance (q<0.1). **d** Sina plots showing the distribution of select V gene copy frequency across TRA and TRB chains in D21 and T21. Points represent individual samples (D21, gray; T21, green); boxes indicate medians and interquartile ranges, with notches approximating 95% confidence intervals. **e-g** Scatter plots showing associations between select B and T cell receptor diversity metrics in individuals with T21. Points represent individual samples with lines indicating fitted linear regression trends with shaded 95% confidence intervals. Panels a-g show TCR-seq of CDR3 regions in T21 (n=131) and D21 (n=60).

Analysis of TCR V and J gene usage revealed differential sequence abundance in DS, with many examples of increased and decreased usage of specific TRA and TRB rearrangements (**Fig. 4j-k**, **Supplementary Fig. 4a-d**). Notably, we observed lower frequency of TRA sequences employed by Mucosal-Associated Invariant T cells (MAITs), such as *TRAV1-2*, *TRAJ20*, and *TRBV6*-family sequences^42,43^ (**Fig. 4j-k, Supplementary Fig. 4a-d**). We previously reported depletion of MAITs from the bloodstream in DS^12,18^, which could be explained by increased migration into inflamed tissues. Likewise, the TCR repertoire skewing in DS suggests a loss of iNKT cells, which predominantly employ *TRAV10* (formally known as Vα24), *TRAJ18*, and *TRBV25-1*^44^, of which *TRAV10* and *TRBV25-1* are decreased in DS (**Fig. 4j-k**). These findings are consistent with previous reports of reduced absolute iNKT cell counts in children with DS^45^. iNKT cells, which primarily recognize lipids in the context of CD1d, have well established roles in protection against pathogens via production of cytokines and cytotoxic granules^46^. Strong evidence from animal models also indicate a role for iNKT cells in controlling autoimmune conditions such as type 1 diabetes (T1D), experimental autoimmune encephalitis, rheumatoid arthritis (RA), and SLE^47^. Thus, reduced iNKT cell frequencies may contribute to impaired control of pathogens and a higher prevalence of autoimmune conditions in DS.

Beyond changes indicative of altered T cell subsets, we also observed TCR changes reported in autoimmune conditions. For example, *TRBV28*, which is significantly enriched in individuals with DS (**Supplementary Fig. 4b**), was found to be associated with both type I diabetes and Crohn’s disease^48,49^. Similarly, the observed reduced usage of *TRBV6-1*, *TRBV6-4*, *TRBV6-5*, *TRBV7-8*, *TRBV14*, and *TRBV16*, along with elevated usage of *TRAV18* in DS is reminiscent of what has been reported in the peripheral CD8 TCR repertoire of patients with vitiligo^50^ (**Fig. 4k, Supplementary Fig. 4a-d**).

We then correlated metrics of diversity across the TCR and BCR datasets. Consistently, individuals with DS with the richest and most diverse TCR repertoires displayed the richest and most diverse BCR repertoires across multiple isotype classes, and those with the strongest usage of the top 5 TCR clones also displayed the strongest usage of the top 5 BCR clones (**Fig. 4l-m**, **Supplementary Fig. 4e-g**).

Altogether, these findings indicated parallel dysregulation of T cell memory tracking with dysregulated humoral immunity in DS.

### Profound disorganization of secondary lymphoid tissue in Down syndrome

Next, we investigated the effects of T21 on secondary lymphoid tissue architecture critical for lymphocyte maturation and adaptive immunity. We conducted a histological and spatial transcriptomic analysis of tonsillar tissue obtained from a cohort of children with and without DS undergoing adenotonsillectomy for treatment of obstructive sleep apnea (OSA). Despite similar age and BMI, tonsils from children with DS displayed significantly reduced tonsil weight compared to controls (**Supplementary Fig. 5a**). Histological examination revealed pronounced structural disorganization and increased interstitial fibrosis in tonsils from children with DS (**Fig. 5a-b**). Hematoxylin and eosin (H+E) staining showed significant reduction in both the number and area of germinal centers in DS (**Fig. 5a**). Masson trichrome staining confirmed increased collagen area across the tonsillar parenchyma, including within germinal centers and mantle zone (MZ), yielding elevated fibrotic scores in the perisinoidal and subepithelial compartments (**Fig. 5b, Supplementary Fig. 5b-c**). Immunofluorescence for CD20 (B cells), CD3 (T cells), and podoplanin (PDPN, a marker for stroma) further demonstrated disorganization of tonsillar architecture, characterized by less-defined germinal centers, elevated CD3 integrated density, and increased staining for PDPN within germinal centers (**Fig. 5c, Supplementary Fig. 5d**).

**Fig. 5.**
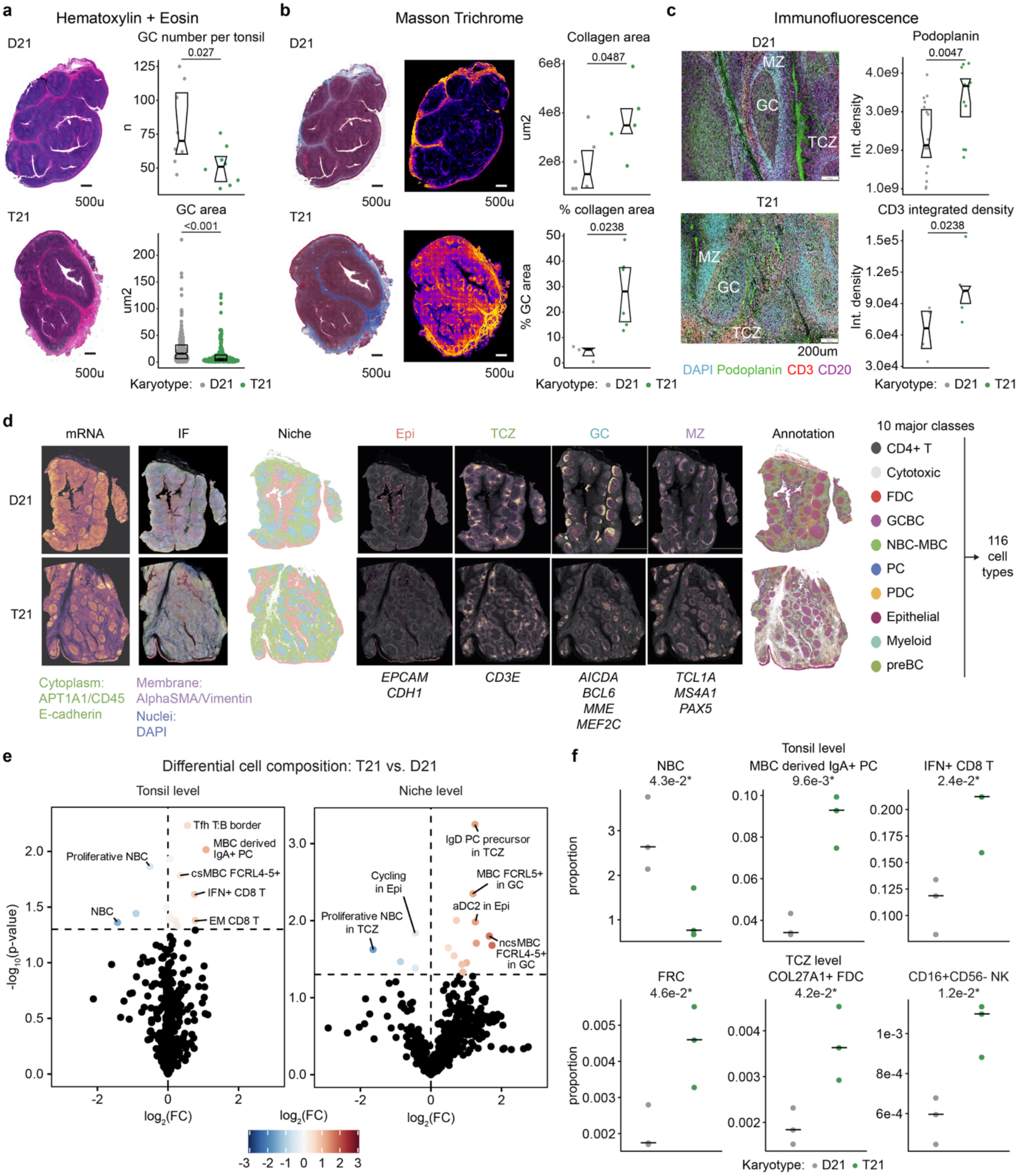
| Profound dysregulation of secondary lymphoid tissue in Down syndrome. **a** Hematoxylin and eosin (H&E) stain of tonsil from donors that are either euploid controls (D21) or with trisomy 21 (T21). Nuclei (blue) and cytoplasm/extracellular matrix (pink). Sina plots display quantification of number of germinal centers (GC, top) and area of GCs (bottom) in tonsils (n=7 D21, n=8 T21). **b** Masson Trichrome (left panel) stain of tonsil depicting collagen (blue), nuclei (black), and cytoplasm (red). Collagen signal (right panel) after color deconvolution. Sina plots display quantification of collagen area and % of GC area containing collagen (n=6 D21, n=5 T21). Scale bars: 500 μm. **c** Overlay of fluorescence images obtained from one field of view (FOV) depicting GC, Mantle Zone (MZ) and T Cell Zone (TCZ) of D21 or T21 tonsils. Green: podoplanin, red: CD3, pink: CD20, and cyan: DAPI. Scale bars: 200 um. Sina plots showing the integrated density of podoplanin per FOV (n=5 donors) and average (n=2 FOV) integrated density for CD3 (n=5 donors). **d** Xenium Spatial transcriptomics work-flow. From left to right: Morphology and mRNA – cell segmentation markers in Xenium Explorer 4.0., where nuclei are blue (DAPI), cytoplasm is green (APT1A1/CD45/E-Cadherin), interior RNA is yellow (18S), and membrane is pink (Alpha SMA/Vimentin); mRNA: density map of total mRNA transcript detected by 10x Genomics Human 5k pan tissue and pathways panel; Niche: niche assignment by Robust Cell Type Deconvolution (RCTD), Epi: Epithelium, TCZ: T Cell Zone, GC: Germinal Center, MZ: Mantle Zone; Annotation: RCTD Spatial Map of high-level cell cluster annotation. mRNA markers listed below niche-specific localization images were used to discriminate individual niches. **e** Volcano plots displaying differential cell composition at the tonsil (left) and niche-specific (right) level. Niche-specific plot includes all four identified tonsil niches (i.e., Epi, TCZ, GC, MZ). Points represent individual cell types with color indicating log2(fold-change), black points indicate non-significant (p≥0.05). **f** Plots displaying select differential frequencies of indicated cell population at the tonsil (top) and TCZ (bottom) niche level. Differences between T21 and D21 for histology and immunofluorescence staining (a-c) were determined by two-sided Mann Whitney U test. Differences in tonsil and niche cell composition (e-f) between T21 and D21 were determined by student’s t-test (n=3).

**Supplementary Fig. 5.**
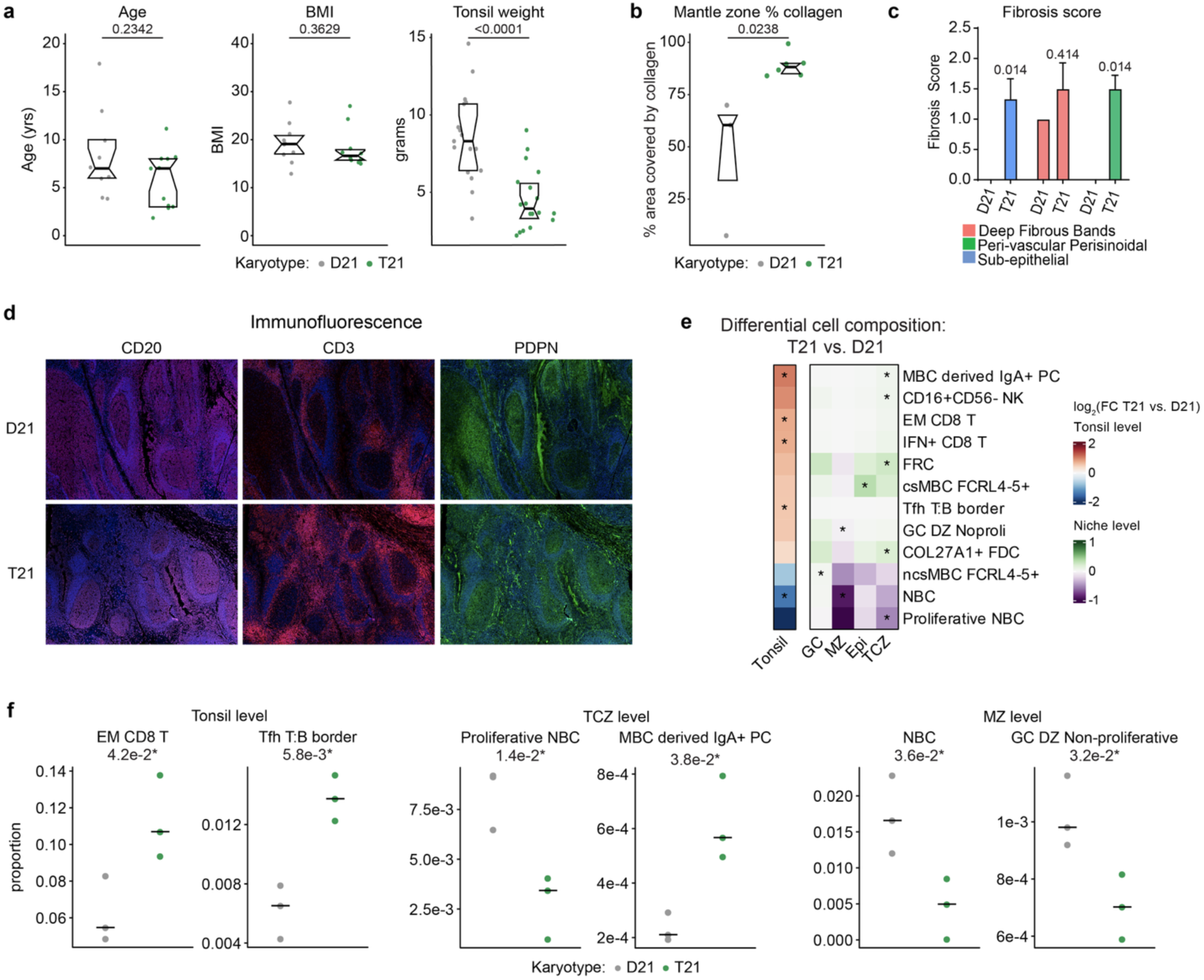
| Strong dysregulation of tonsil architecture in Down syndrome. **a** Sina plots showing the age, BMI, and tonsil weight in donors that are either euploid controls (D21) or individuals with trisomy 21 (T21). Statistics from Mann Whitney U test. **b** Sina plot displaying % of mantle zone area containing collagen. **c** Bar chart of blinded pathologist scoring for tonsil fibrosis (n=6 T21,n=4 D21) and analyzed by Mann Whitney U test with Holm Sidak method to correct for multiple comparisons. Stats above bars indicate adjusted p-values. **d** Representative images of CD20 + DAPI, CD3 + DAPI, and Podoplanin + DAPI. **e** Heatmap displaying differential cell composition at the tonsil (left) and niche-specific (right) level. Differences were determined by Student’s t-test with asterisks indicating significance (p<0.05). **f** Plots displaying select differential frequencies of indicated cell populations at the tonsil (left), TCZ niche (middle) and MZ niche (right) level.

Prompted by these observations, we then performed spatial transcriptome analysis using the Xenium platform (measuring ∼5,000 mRNAs) in conjunction with Robust Cell Type Deconvolution^51^ (see **Methods**). We mapped the spatial landscape across four distinct niches: the epithelium (Epi), the MZ, germinal centers, and the T cell zone (TCZ), and deconvoluted 10 major high-level clusters with subsequent low-level annotation of 116 unique cell types/states (**Fig. 5d, Supplementary Table 5**). Differential analysis of cell type proportions revealed profound tonsil-level and niche-specific alterations in DS (**Fig. 5e-f**, **Supplementary Fig. 5e-f**). Naive B cells (NBCs) and proliferative NBCs are depleted within the entire tonsil and within the MZ and TCZ. The memory B cell (MBC) compartment is shifted toward MBC-derived IgA+ plasma cells across the entire tonsil and within the TCZ, consistent with increased IgA isotype usage in DS. We also observed increased frequencies of atypical FCRL4+ and FCRL5+ MBCs, which have been found to be associated with chronic inflammatory states, rheumatoid arthritis, and multiple sclerosis^52,53^. These changes in B cell subsets are mirrored by increased frequencies in IFN+ CD8+ T cells and effector memory (EM) CD8+ T cells across the entire tonsil, indicative of T cell activation and differentiation, along with increased frequencies of Tfh T:B border cells, a transitional population of activated CD4+ T cells located at the T cell-B cell interface (**Fig. 5f, Supplementary Fig. 5e-f**). Alterations in lymphoid lineages are accompanied by multiple changes in mesenchymal stromal cells and innate immune cell types in the TCZ, such as higher proportions of fibroblastic reticular cells (FRCs), follicular dendritic cells (FDCs) expressing COL27A1, and CD16+ CD56-NK cells (**Fig. 5f, Supplementary Fig. 5e-f**).

Together, these observations demonstrate clear disorganization of secondary lymphoid tissue architecture and niche composition in DS, providing a structural basis for peripheral adaptive immune dysregulation.

### Transcriptome changes underlying disrupted adaptive immune environment in Down syndrome

To characterize the gene expression changes underlying the structural remodeling and shifts in cell composition observed in tonsils of children with DS, we completed differential gene expression analysis across all resolved cell subsets within the parent clusters, with emphasis on cell types with the highest number of differentially expressed genes (DEGs) and the DEGs which were consistently changed across multiple subsets (**Fig. 6a, Supplementary Fig. 6a-b, Supplemental Table 6**).

**Fig. 6.**
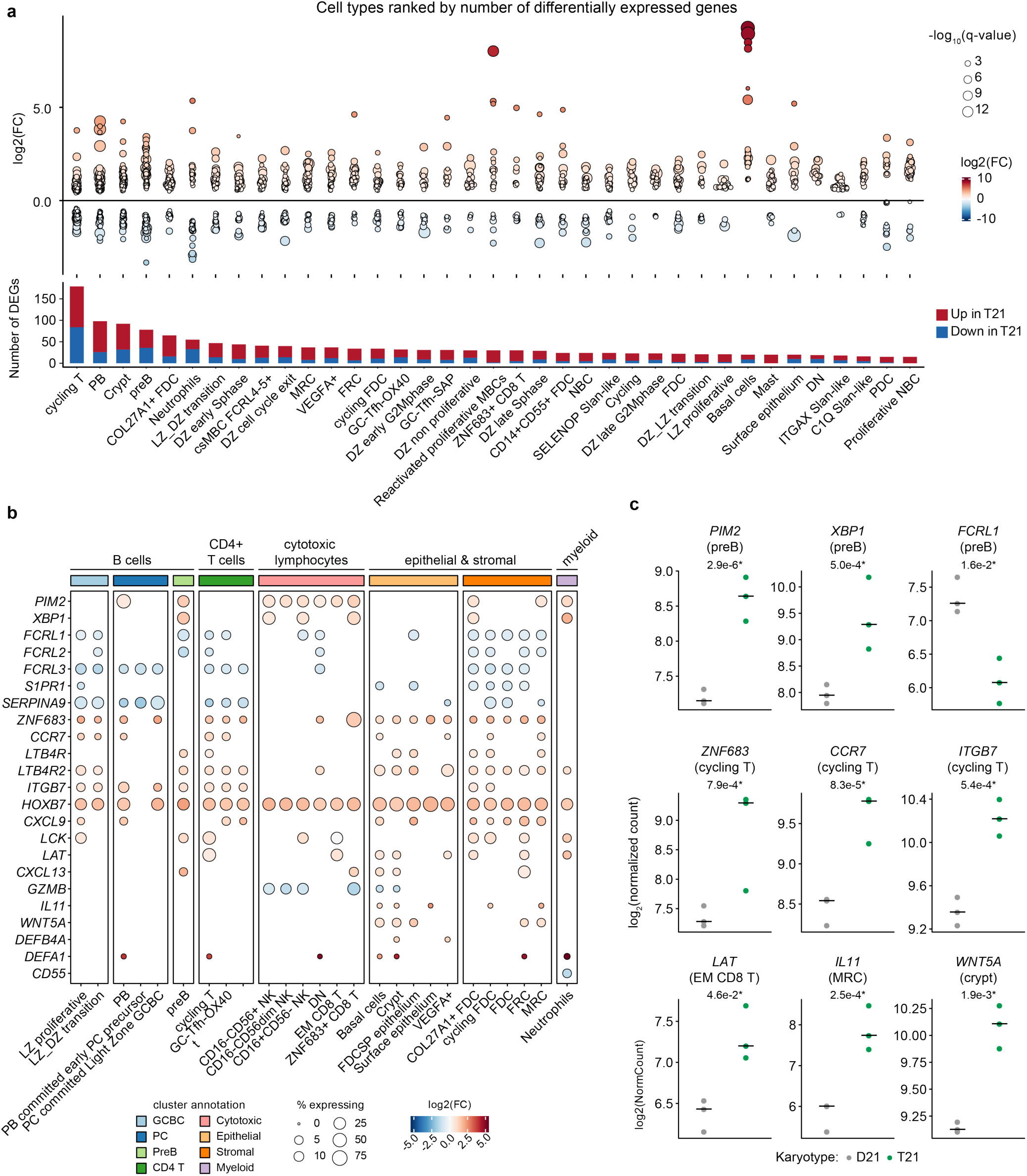
| Gene expression changes reveal highly disrupted adaptive immune environment in Down syndrome. **a** Differential gene expression from pseudo-bulk RNA sequencing of unique tonsil cell populations comparing individuals with trisomy 21 (T21, n=3) vs. euploid controls (D21, n=3). Differential expression was analyzed by DESeq2. Multiple hypothesis correction was performed using the Benjamini-Hochberg method. Selected cell types include those with at least 15 differentially expressed genes. Sina plots (top) display the distribution of differentially expressed genes per cell type, with color indicating log2(fold-change) and size corresponding to –log10(q-value). Bar graphs (bottom) indicate the number of upregulated (red) and downregulated (blue) genes per cell type. **b** Bubble plot displaying differential expression of select genes across select tonsil cell populations grouped into major cellular compartments. Dot size indicates the percentage of cells expressing each gene, and dot color indicates differential expression (log2 fold change). Sample sizes as in (a). **c** Representative normalized gene expression in selected cell populations in D21 and T21. Each point represents an individual sample, with horizontal bars denoting the group medians. Sample sizes as in (a).

**Supplementary Fig. 6.**
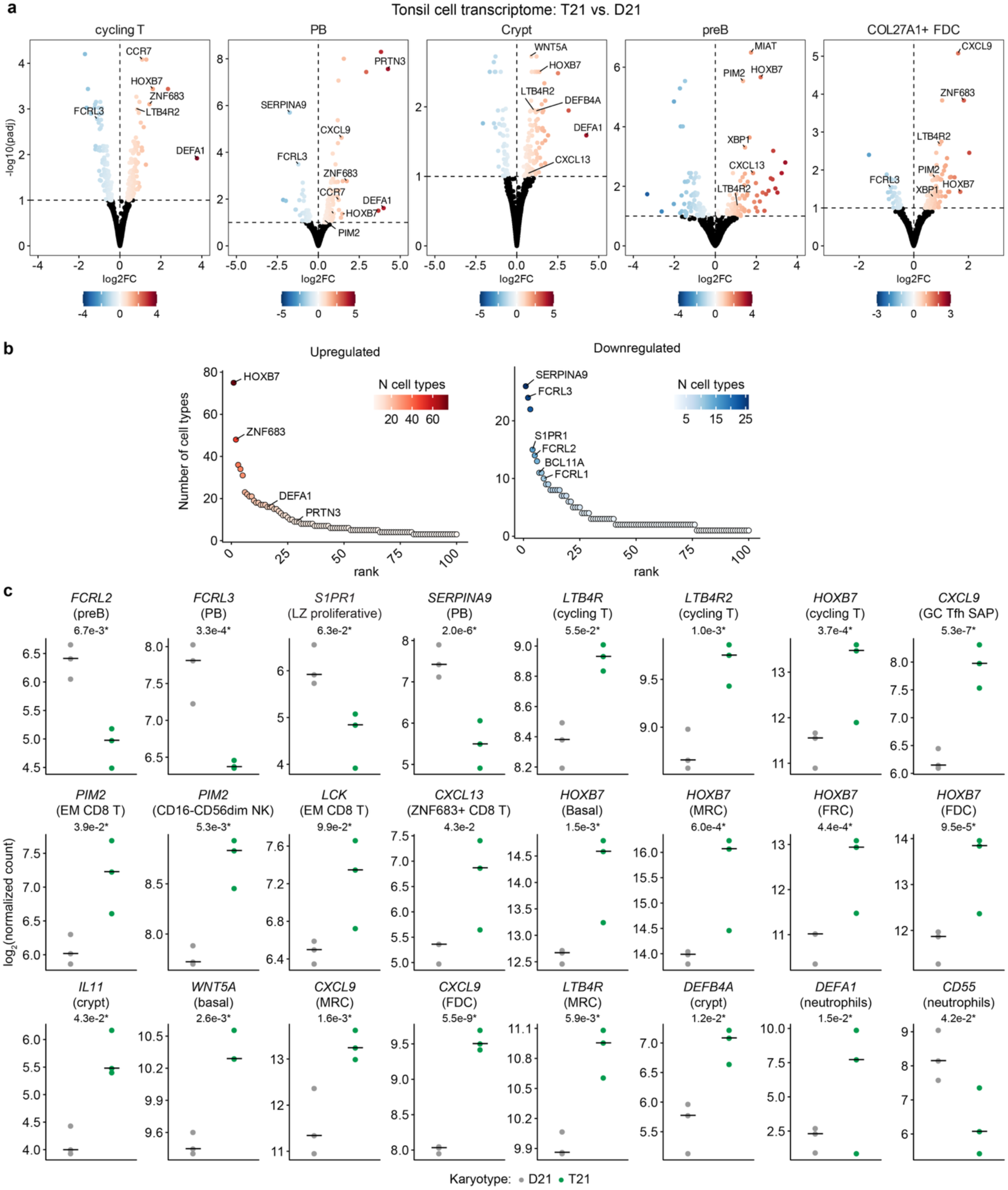
| Gene expression changes reveal highly disrupted adaptive immune environment in Down syndrome. **a** Volcano plots displaying differential gene expression from pseudo-bulk RNA sequencing of select tonsil cell populations comparing individuals with trisomy 21 (T21, n=3) vs. euploid controls (D21, n=3). Differential expression was analyzed by DESeq2. Multiple hypothesis correction was performed using the Benjamini-Hochberg method. Points are colored by log2(fold-change) with black points representing non-significant differences (q≥0.1). **b** Rank plots showing the 100 genes most frequently upregulated (left) or downregulated (right) across cell populations. Genes were ranked by the number of cell populations in which they were significantly differentially expressed, with color indicating the number of cell populations exhibiting differential expression. **c** Representative normalized gene expression in selected cell populations in D21 and T21. Each point represents an individual sample, with horizontal bars denoting the group medians. Sample sizes as in (a).

Tonsillar B cell populations from children with DS show signatures of increased differentiation toward plasmablasts and plasma cells, including increased expression of *PIM2,* a protein kinase upregulated upon terminal differentiation of B cells^54^, and X-Box Binding Protein 1 (*XBP1*), a transcription factor crucial for plasmablast differentiation and antibody production^55^ (**Fig. 6b-c**). We also observed reduced expression of Fc Receptor Like 1, 2, and 3 (*FCRL1-3*), a group of immunoglobulin superfamily receptors downregulated during plasma cell differentiation^56^, and *S1PR1*, a lymphoid egress receptor downregulated in resident plasma cells^57^ (**Fig. 6b-c, Supplementary Fig. 6b**). Notably, *SERPINA9*, a canonical marker of germinal center B cells, is the most consistently downregulated gene across multiple cell types (**Supplementary Fig. 6b-c**), consistent with the decreased number and area of germinal centers.

Among tonsillar CD4+ T cell subsets, we observed changes indicative of dysregulated activation, homing, and tissue residence in DS, including upregulation of *ZNF683* (Hobit), *CCR7*, the leukotriene B4 (LTB4) receptors *LTB4R* and *LTB4R2*, *ITGB7*, and *HOXB7* (**Fig. 6b-c, Supplementary Fig. 6b**). ZNF683 is a transcription factor that promotes tissue adaptation and long-term persistence of antigen-experienced CD4+ T cells^58^. CCR7 serves as a lymphoid homing receptor for T cells, guiding naïve and memory cells into lymphoid tissues via CCL19 and CCL21 gradients^59^. LTB4R expression is highly induced in both CD4+ and CD8+ T cells upon TCR activation and mediates the potent chemotactic effects of LTB4^60^. ITGB7 is an integrin involved in lymphocyte homing to lymphoid and epithelial tissues^61^. HOXB7 is a developmental transcription factor reactivated in pathological contexts known to drive ECM remodeling, EMT, and fibrosis^62^. Overexpression of these homing and residence markers within cycling T cells suggests local retention and expansion of these tissue-adapted lymphocytes. Additionally, tonsillar CD4+ T cells from children with DS overexpress *CXCL9*, an IFN-inducible chemokine that recruits CXCR3-expressing activated T cells^63^. Overexpression of *CXCL9* was also observed across many other tonsillar cell types, consistent with hyperactive IFN signaling in DS (**Fig. 6b-c, Supplementary Fig. 6b**).

Cytotoxic lymphocyte populations display changes indicative of persistent activation and altered effector functions in DS (**Fig. 6b-c, Supplementary Fig. 6b**). EM CD8 T cells exhibit increased expression of *PIM2*, which is induced by cytokine signaling and TCR stimulation^64^. *PIM2* is also elevated in NK cells and ILC1s, consistent with a shared program of chronic activation. Additional signs of increased T cell activation include upregulation of *LCK* and *LAT*, two key mediators of TCR downstream signaling^65^. Notably, the ZNF683+ CD8 T population also exhibited increased expression of *CXCL13*, a chemokine that plays a central role in B cell recruitment^66^.

Within the epithelial and stromal compartments, we observed changes consistent with the marked fibrotic phenotype observed histologically (**Fig. 6b-c, Supplementary Fig. 6b**). Across epithelial cells, FRCs, FDCs, and marginal reticular cells (MRCs), one of the most consistent transcriptional changes is upregulation of *HOXB7*, a master regulator of tissue remodeling^62^. Furthermore, we observed increased expression of other genes involved in ECM remodeling, including *IL11,* a major driver of fibrosis in a number of pathological contexts^67^, and *WNT5A*, which activates fibroblasts to drive production of ECM components^68^. WNT5A also decreases B cell proliferation through the non-canonical Wnt pathway via downregulation of cyclin D1^69^. Additionally, epithelial and stromal populations show signs of a pro-inflammatory phenotype, including robust overexpression of *CXCL9* and LTB4 receptors^70^.

Lastly, the transcriptome of T21 myeloid cells in the tonsil shows alterations indicative of dysregulated myeloid cell activation, such as elevated expression of *DEFA1,* an antimicrobial peptide and chemoattract driving mobilization of naïve T cells and immature dendritic cells^71^, along with decreased expression of *CD55,* a negative regulator of neutrophil activation that suppresses complement activation by preventing assembly of the C3 convertase^72^ (**Supplementary Fig. 6c**).

Together, these results show that T21 is associated with drastic changes in the transcriptome of tonsillar tissue indicative of a highly dysregulated lymphocyte maturation environment.

## DISCUSSION

Over the past several decades, advances in social inclusion and medical care have dramatically increased life expectancy for individuals with DS. However, further gains in both longevity and quality of life are increasingly constrained by early-onset Alzheimer’s disease and persistent immune dysregulation, which is now recognized as a core feature of DS^13^. Individuals with DS exhibit increased rates of autoimmune diseases, impaired vaccine responses, severe complications from infections, and a state of low-grade systemic inflammation that emerges early in life and persists across the lifespan, likely contributing to precocious aging and frailty^2,6,9,12,13^. As traditional causes of early mortality have become more manageable, immune dysregulation has emerged as a major biological barrier limiting healthy aging in DS. Therefore, a deeper understanding of the mechanisms driving immune dysfunction, and the development of targeted immunomodulatory interventions, represent critical opportunities to further improve quality of life and increase life expectancy in this growing population. Within this context, we report here an extensive investigation of adaptive immunity in DS, which produced many interesting observations relative to earlier studies.

Our lifespan analysis reveals that T21 exerts dynamic effects on the B cell lineage, beginning with signs of impaired adaptive immune memory in childhood and progressing toward increasingly activated, inflammatory, and autoreactive states during adulthood. Early depletion of non-switched memory B cell populations suggests defects in germinal center output and maturation, providing a plausible explanation for the increased susceptibility to infection and suboptimal vaccine responses. This is followed by progressive expansion of CD11c+, T-bet+, CD95+ switched memory B cell populations, ABCs, and plasmablasts, cellular states linked to persistent immune stimulation, autoimmunity, and immunosenescence^73^. The close correlation between these phenotypic shifts and lifelong elevation of basal STAT1 phosphorylation suggests that chronic activation of the IFN/JAK/STAT axis may serve as a central driver of B cell remodeling in DS. Notably, the temporal trajectory observed here mirrors patterns described in chronic inflammatory and autoimmune disorders, where sustained immune activation progressively reshapes the B cell compartment away from diverse, flexible memory responses and toward terminally differentiated and autoreactive states^74,75^. These findings also support a model in which T21 accelerates immune aging across the B cell lineage, resulting in the simultaneous coexistence of immunodeficiency, impaired immune memory, and increased autoreactivity.

A notable finding of this study is the substantial inter-individual heterogeneity in B cell depletion observed in DS within an overall acceleration of age-dependent B cell loss. By leveraging this heterogeneity, we identified a subgroup of individuals with particularly severe B cell lymphopenia that exhibited widespread immune remodeling extending well beyond the B cell lineage and heightened IFN transcriptional signatures. These findings suggest that B cell lymphopenia is not merely a consequence of impaired lymphopoiesis but rather a marker of a broader inflammatory state associated with chronic immune activation. Importantly, elevated expression of IFNRs in those with severe B cell lymphopenia further supports a model in which gene dosage effects sensitize immune cells to IFN signaling, resulting in chronic pathway activation and progressive depletion of naïve lymphocyte pools, along with expansion of activated and antibody-producing immune cell states.

Our analyses of the BCR and TCR repertoires demonstrated progressive restriction and skewing across the lifespan in DS. Consistently, the changes observed mimic findings obtained during the study of adaptive immunity in chronic infections and/or autoimmune diseases^33–35^. In healthy individuals, broad and diverse antigen receptor repertoires enable effective responses to a wide range of pathogens while maintaining tolerance to self-antigens. In contrast, many autoimmune conditions exhibit repertoire impoverishment, characterized by reduced diversity, diminished richness, and contraction of the naïve lymphocyte pool, together with repertoire skewing, reflected by overrepresentation of specific V(D)J gene segments, clonal expansions, and altered CDR3 characteristics^34^, all of which is now also evident in DS. The observation of parallel BCR and TCR repertoire abnormalities provides evidence for a systemic breakdown of adaptive immune homeostasis and supports the concept that autoimmune pathology in DS arises from coordinated dysregulation of both humoral and cellular immunity rather than defects confined to a single lymphocyte lineage^76^.

Our analysis of tonsillar tissue reveals that immune dysregulation in DS extends beyond circulating immune cells and is rooted, at least in part, in profound abnormalities of secondary lymphoid tissue architecture. The marked reduction in germinal center number and size, together with extensive fibrosis, stromal remodeling, and disruption of normal T cell and B cell compartmentalization, suggests that T21 fundamentally alters the tissue niches required for adaptive immune education^77^. These structural abnormalities were accompanied by depletion of naïve B cells, expansion of atypical FCRL4+ and FCRL5+ memory B cells associated with chronic inflammatory states^52,53^, and accumulation of activated cytotoxic T cell populations. Such changes are consistent with a model in which chronic IFN signaling and persistent immune activation progressively remodel secondary lymphoid organs, impairing the generation of balanced humoral immunity while favoring the emergence of activated and potentially pathogenic lymphocyte states^78^. The concomitant expansion of FRCs, FDCs, and collagen-rich stromal networks further suggests that aberrant tissue repair and fibrosis may contribute to the loss of normal lymphoid organization^79,80^. Together, these findings provide a mechanistic link between T21, altered lymphoid tissue microenvironments, and the widespread defects in B cell memory and T cell differentiation observed in DS.

Our spatial transcriptomic analyses demonstrate coordinated transcriptional changes across virtually every cellular compartment, supporting a model in which chronic inflammatory signaling, tissue remodeling, and disrupted lymphocyte differentiation reinforce one another to impair germinal center function. Within the B cell compartment, increased expression of *PIM2* and *XBP1*, together with reduced *FCRL1-3*, *S1PR1*, and S*ERPINA9*, is consistent with a shift from germinal center maintenance toward terminal plasma cell differentiation. Concurrently, CD4+ and cytotoxic lymphocytes exhibit transcriptional signatures of persistent activation, tissue adaptation, and altered migratory behavior, accompanied by widespread overexpression of the IFN-inducible chemokine CXCL9, supporting a central role for chronic IFN signaling in organizing the inflammatory milieu. Finally, coordinated upregulation of *HOXB7*, *IL11*, and *WNT5A* across epithelial and stromal compartments provides a plausible molecular basis for the extensive fibrosis and ECM remodeling observed. Collectively, these findings suggest that T21 fundamentally disrupts the lymphoid microenvironment, shifting secondary lymphoid tissues away from efficient germinal center-dependent immune education and toward a chronically inflamed, fibrotic state that likely contributes to dysregulated adaptive immunity DS.

This study has several limitations. First, our report does not describe associations between the cellular and molecular events observed and various developmental and/or clinical characteristics of DS, which would require a significant effort toward rich annotation of clinical metadata and health outcomes for the hundreds of research participants that contributed blood samples to this study. Future studies would be required to define phenotypic associations. Second, given limited sample availability, we could not complete all assays on all samples. Whereas some blood samples were concurrently analyzed for generation of multiple datasets, which enabled interesting cross-platform comparisons, this was not feasible across the full sample set. Third, caution should be exercised in the interpretation of the associations described here when considering cause-effect relationships. Although some associations are suggestive of causal relationship (e.g., increased IFN/JAK/STAT signaling as a driver of B cell dysregulation), mechanistic studies in experimental models would be required for full ascertainment of directionality. Fourth, because sampling of pediatric tonsil tissue is ethically unfeasible outside the context of surgical procedures to alleviate OSA, our analysis could be limited by the potential divergent etiology of OSA between groups. For example, in euploid controls, pediatric OSA is driven by primary lymphoid tissue hypertrophy, whereas in DS it could stem primarily from craniofacial anomalies and pharyngeal hypotonia, and these differences could affect interpretation of our comparisons.

In sum, the observations reported here significantly expand our understanding of immune dysregulation in DS, while also emphasizing the pursuit of immunomodulatory therapies for improved health outcomes and further gains in life expectancy in this population.

## METHODS

### Study participants

The analyses from blood samples presented in this manuscript are part of an ongoing study of the population with DS at the Linda Crnic Institute for Down Syndrome known as the Human Trisome Project (HTP, NCT02864108, www.trisome.org). Data from the HTP has been previously published in other contexts^4,5,18,81,82^. HTP participants were enrolled under a protocol approved by the Colorado Multiple Institutional Review Board (COMIRB 15-2170), with all participants or their legal guardians providing written informed consent. The biological datasets analyzed were generated from de-identified blood samples and linked to demographic and clinical metadata, with all participants consenting to the sharing of their de-identified information. Tonsil tissues were collected under protocol COMIRB 17-2159. Tonsil samples were classified as medical waste. Tonsil inclusion criteria included clinically indicated extra-capsular tonsillectomy for OSA with no clinical evidence of infection. Sex was defined by sex assigned at birth by self-report.

### Inclusion and ethics

This study included local researchers throughout the entire research process. This research is relevant both locally and globally. This has been determined in collaboration with local advocacy organizations supporting the population with DS. The roles and responsibilities of the authors and collaborators were determined both ahead of research and as needed throughout the process. This research was not restricted or prohibited in the setting where it was conducted. Participants were enrolled under a protocol approved by the Colorado Multiple Institutional Review Board (COMIRB 15-2170, NCT02864108, see also www.trisome.org). While this research does not result in stigmatization, incrimination, discrimination or otherwise personal risk to participants, all information has been de-identified to ensure the safety and wellbeing of research participants. All research activities and experimental procedures were carried out with strict compliance with all federal, state, local and university regulations to mitigate any potential risks to the collaborators. Biospecimens utilized in this research are available through the Human Trisome Project Biobank and can be requested online through www.trisome.org.

### Biospecimen collection and processing

The biological datasets analyzed in this study were derived from peripheral blood samples and matched with demographic and clinical metadata. Peripheral blood was collected using PAXgene RNA Tubes (Qiagen) and BD Vacutainer K2 EDTA tubes (BD). Whole blood from PAXgene tubes was processed for RNA-seq, while samples from EDTA tubes were used for plasma proteomics via SomaScan, as detailed below. Two 0.5 mL aliquots of whole blood from each EDTA tube were also processed for mass cytometry, as described. Remaining EDTA blood samples were centrifuged at 700 x g for 15 minutes to separate plasma, buffy coat (white blood cells), and red blood cells, followed by aliquoting, flash freezing, and storage at –80°C. All centrifugation and storage procedures were completed within two hours of sample collection. Tonsil samples were stored on ice after same day collection via extracapsular tonsillectomy, before Formalin Fixed Paraffin Embedding (FFPE) at the CU Anschutz Gates Institute Histology Core using a VIP5 tissue processor and Histostar embedding unit.

### Whole blood RNA-sequencing (RNA-seq)

RNA was extracted and purified from whole blood collected in PAXgene RNA tubes using the PAXgene Blood RNA Kit (PreAnalytiX). The RNA quality was assessed with a 2200 TapeStation system (Agilent), concentration was measured using a Qubit fluorometer (Invitrogen). Globin RNA was depleted using the GlobinClear kit (ThermoFisher Scientific). Poly-A(+) RNA enrichment and strand-specific library preparation were performed using the NEBNext Poly(A) mRNA Magnetic Isolation Module and the NEBNext Ultra II Directional RNA Library Prep Kit for Illumina (New England Biolabs). Sequencing was conducted by Novogene Co., Ltd on an Illumina NovaSeq 6000 platform, generating paired-end 150 bp reads, with data delivered in FASTQ format.

### SomaScan plasma proteomics

EDTA plasma (125 µL) was analyzed using the SomaScan platform at SomaLogic, Inc., following established proprietary protocols^83^. Briefly, SOMAmer reagents selectively bind target peptides in the sample and levels are quantified on custom Agilent hybridization chips. Data normalization and calibration were performed in accordance with the SomaScan Data Standardization and File Specification Technical Note (SSM-020)^83^. The final output for protein abundance is expressed in relative fluorescent units (RFU).

### Immune cell cytometry by time of flight (CyTOF)

Mass cytometry was performed using two 0.5 mL aliquots of EDTA whole blood. BD Phosflow Lyse/Fix Buffer 5X (BD Biosciences) was used for red blood cell lysis and white blood cell fixation. Following fixation, white blood cells were washed with PBS (Rockland), resuspended in Cell Staining Buffer (Fluidigm), and stored at −80°C. Prior to staining, samples were thawed at room temperature, washed with Cell Staining Buffer, and barcoded using the Cell-ID 20-Plex Pd Barcoding Kit (Fluidigm). Barcoded samples were pooled into batches of up to 19 samples, with each batch including a shared reference sample. Antibodies used for staining were either pre-conjugated to metal isotopes or conjugated in-house using the Maxpar Antibody Labeling Kit (Fluidigm; details provided in **Supplementary Table 1**). Working antibody dilutions were titrated and validated using the reference sample and benchmarked against relative frequencies obtained through independent flow cytometry. Surface marker staining was performed at 4°C for 30 minutes in Cell Staining Buffer with Fc Receptor Binding Inhibitor (eBioscience/ThermoFisher Scientific), followed by washing. Cells were then permeabilized with Buffer III (BD Pharmingen Transcription Factor Phospho Buffer Set) at 4°C for 20 minutes, then washed with perm/wash buffer from the same kit. Intracellular staining for transcription factors and phospho-epitopes was conducted at 4°C for 1 hour in perm/wash buffer, followed by a final wash in Cell Staining Buffer. Barcoded and stained cells were labeled with Cell-ID Intercalator-Ir (Fluidigm) before analysis on a Helios mass cytometer (Fluidigm). Data were exported as FCS v3.0 files for downstream pre-processing and analysis.

### B cell and T cell receptor (BCR/TCR) library preparation and sequencing

Whole blood RNA was used for immune receptor sequencing. B cell receptor (BCR) heavy-chain and T cell receptor (TCR) libraries were prepared from RNA using the TaKaRa SMART-Seq Human BCR and TCR kits, respectively, on an automated liquid handling system (epMotion 5070), following the manufacturer’s protocols. These approaches use reverse transcription and targeted amplification of immunoglobulin– and TCR-encoding transcripts, thereby restricting sequencing to receptor-derived cDNA rather than genomic DNA or the full transcriptome. Prepared libraries were barcoded, pooled, and normalized prior to sequencing. Library quality control and quantification were performed using a QuantStudio 7 system (Applied Biosystems) to assess amplifiable library concentration and confirm the presence of appropriate sequencing adapters. Libraries were normalized to approximately 2 nM prior to sequencing. Sequencing was performed on an Illumina NextSeq 2000 platform (P2 flow cell configuration), generating approximately 5 million reads per sample.

### Tonsil histology and immunofluorescence

Paraffin blocks were microtome sectioned at a depth of 5 micron and stained for Hematoxylin and Eosin (H&E) and Masson Trichrome at the Gates Institute Histology Core at CU Anschutz. Following Antigen Retrieval in DIVA Decloaker, tonsil sections were stained with anti-CD3, anti-Podoplanin, anti-CD20, goat anti-IGg2b AF555 goat anti-Rabbit DyLight 755, goat anti-Rat 488 and DAPI and mounted in Prolong Glass anti-fade. Slides were imaged at 20x magnification on an Olympus VS200 Slide Scanner at the ALMC core, and images were exported as .vsi files for analysis.

### Spatial transcriptomics of tonsils

FFPE tonsil sections were placed within the printable area of a Xenium slide and sample preparation, probe hybridization, ligation and amplification were carried out using the Xenium Prime In situ gene expression kit (Xenium Prime 5000K Human Pan Tissue and Pathways Assay Kit,10X Genomics), according to the guidelines laid out in 10X Genomics’ Demonstrated Protocols CG000578, CG000580, and CG000749. Multimodal segmentation for precise cell boundary delineation was performed using the Xenium Cell Segmentation Staining Reagents (10x Genomics), containing DAPI and a cocktail of membrane/cytoplasmic protein markers (Alpha SMA/Vimentin and 18SRNA/ATP1A1/CD45/E-Cadherin). Images were acquired on the Xenium Analyzer, and transcript decoding was executed using the Xenium Onboard Analysis Pipeline.

### Image analysis

All image analysis was carried out in FIJI (v. 1.54f https://fiji.sc). Collagen area was measured by applying a Binary Collagen Mask to the unprocessed image. The mask was constructed by Masson Trichrome Dye Vector deconvolution with ColourDeconvolution2, followed by background subtraction (Rolling Ball Sliding Paraboloid 100 pixel), enhance local contrast (Block Size 56, Slope 4), median filtering (2 pixel) and Otsu thresholding on the deconvoluted blue channel. Number and area of germinal centers were measured by delineation of germinal centers as individual Regions of Interest (ROI) per tonsil section under pathologist guidance and measured using analyze particle function. CD3 and podoplanin integrated density were measured on the raw image using an Otsu-Threshold Binary Mask, which was pre-filtered using Top-Hat (30 pixel or 50 pixel) and gaussian blur (sigma=1). Analyze particle function was performed at a minimum size of 2 um to remove small noise artifacts.

### Statistical analysis

#### Data analysis and visualization

Data pre-processing, statistical analyses, and visualization were conducted in R (R 4.0+, RStudio 2022+, Bioconductor 3.16+), as detailed below. For all analyses, multiple hypothesis correction was performed using the Benjamini-Hochberg method for each family of tests. Plots were generated primarily using ggplot2 (v3.4.4). Sina plots comparing distributions between groups, were created with the ggplot2 and the *geom_sina()* function from ggforce (v0.4.1), with points horizontally jittered based on local density and medians and interquartile ranges displayed with overlaid boxplots. Heatmaps were generated using the tidyheatmap (v1.8.1) package. Color palettes were sourced from RColorBrewer (v1.1.3).

#### Analysis of immune cell CyTOF data

*Pre-processing.* Bead-based normalization and removal was carried out using the Matlab-based Normalizer tool (version 9.12). Batched FCS files were demultiplexed using the Matlab-based Single Cell Debarcoder tool^84^. Reference-based normalization of individual samples across batches was then carried out using the R script *BatchAdjust()*. Per sample FCS files were then uploaded to CellEngine (CellCarta, version accessed 2021) and gated on live singlets (i.e., PARP negative, excluding CD3+CD19+ doublets). These live singlets were exported as individual FCS files per sample. Next, hematopoietic lineage (CD45-positive) non-granulocytic (CD66-low) cells were gated and exported. Finally, CD3-positive and CD19-positive cells were exported as ‘T cells’ and ‘B cells’, respectively. Each set of FCS files was then subsampled to a maximum of 50,000 events per sample (for live cells and CD45+CD66low, T cells) or all B cells available for subsequent analysis in R.

*Unsupervised clustering.* For each parent population above, all 388 per-sample FCS files were imported into R as a flowSet object using *read.flowSet()* function from the flowCore package. Next, a SingleCellExperiment object was constructed from the flowSet object using the *prepData()* function from the CATALYST package, applying Arcsinh transformation with cofactors optimized using flowVS package. Quality control and diagnostic plots were examined with the help of functions from the CATALYST and tidySingleCellExperiment packages^85^. Unsupervised clustering using the FlowSOM algorithm^86^ was carried out using the *cluster()* function from the CATALYST package, with grid size set to 10 x 10 to give 100 initial clusters and a maxK value of up to 40 was explored for the subsequent meta-clustering.

*Cell type annotation.* To aid in assignment of clusters to specific lineages and cell types, the MEM package (marker enrichment modeling) was used to call positive and negative markers for each cell cluster based on marker expression distributions across clusters. Manual review and comparison to marker expression histograms allowed for high-confidence assignment of most clusters to specific cell types. Relative frequencies for each cell type / cluster were then calculated for each sample as a percentage of the total of the parent population (Live, CD45+CD66low, T cells, B cells).

*Linear model analysis*. To identify markers with differential expression in B cells, we used linear regression. Extreme outliers (*Q1 – 3IQR, Q3 + 3IQR*) on a per-group and per-marker basis were excluded from further analysis. Differential expression analysis was carried out in R by linear regression using the *lm()* function from the stats package (v4.4.0). For comparisons between individuals with T21 and D21, asinh-transformed mean expression served as the outcome/predictor variable and T21 status served as the predictor/independent variable, with age, sex and sample source as covariates. For visualization purposes only, *removeBatchEffect()* from the limma (v3.60.4) package was used to adjust for these same covariates.

*Beta regression analysis.* To identify differential abundance among B cell types and to determine differential pSTAT1 positivity in B cell types, we used beta regression. Extreme outliers (*Q1 – 3IQR, Q3 + 3IQR*) on a per-group and per-cell type basis were excluded from further analysis. Beta regression analysis was carried out using the *betareg* package (v3.1-4) in R. For each model, either cell type relative frequencies or the proportion of pSTAT1-positive cells within each cell type served as the outcome/dependent variable and either T21 status or B cell low/high subgroup served as the independent/predictor variable, with adjustment for age, sex, and sample source. Correction for multiple comparisons was performed using the Benjamini-Hochberg method. For visualization purposes only, *adjust()* from the datawizard (v1.3.1) package was used to adjust for these same covariates.

*Differential Expression Sliding Window Analysis (DESWAN)*. DESWAN was used to evaluate the effects of T21 status across the lifespan using DEswan R package (v0.0.0.9001). Analyses were conducted within overlapping 10-year windows, sliding incrementally across the lifespan from a center of 3 years to a center of 57 years. Analyses were restricted to this range to ensure windows with a minimum of three samples per group. Differential abundance within each window was determined using methods specific to each data type on data pre-adjusted for technical effects, specifically sample source, using adjust() from datawizard (v1.3.1) package. Differences related to T21 status were determined by comparing individuals with T21 to those with D21 within each window, using age group and sex as covariates. Multiple hypothesis correction was performed in each window using the Benjamini-Hochberg method, with a significance threshold of q < 0.1, corresponding to a false discovery rate (FDR) of 10%.

*Spearman correlation*. Spearman correlation coefficients (rho) and p values were calculated for the B cell marker expression in T21 using the *rcorr*() function from the Hmisc package (v 4.4-0). Correlations were performed on data pre-adjusted for age, sex, and sample source, using adjust() from datawizard (v1.3.1) package. Multiple testing correction was performed separately for each marker using the Benjamini-Hochberg method, with a significance threshold of q < 0.1, corresponding to a false discovery rate (FDR) of 10%.

*B cell high and low groups.* B cell frequencies were adjusted for age, sex, and sample source using *adjust()* from the datawizard package (v1.3.1). A local polynomial regression (LOESS) fit was generated in the T21 group using the *loess()* function in R. Residuals from the fitted curve were calculated for each individual, and those with residuals in the upper and lower 25^th^ percentiles were classified as the B cell high and B cell low groups, respectively.

#### Analysis of whole blood RNA-seq data

FASTQ files generated by Novogene provided RNA-seq data yielding ∼33-103×10^6^ raw reads and ∼21-69×10^6^ final mapped reads per sample. Data quality was assessed using FASTQC (v0.11.5) and FastQ Screen (v0.11.0). Low-quality reads were trimmed and filtered using bbduk from BBTools (v37.99) and fastq-mcf from ea-utils (v1.05). Low-quality reads were trimmed and filtered using bbduk from BBTools (v37.99) and fastq-mcf from ea-utils (v1.05). Reads were aligned to the GRCh38 human reference genome using HISAT2 (v2.1.0) in in paired spliced-alignment mode, with a GRCh38 index and Gencode v33 basic annotation GTF serving as references. Alignments were sorted and filtered for mapping quality (MAPQ > 10) using Samtools (v1.5). Gene-level count data were quantified using HTSeq-count (v0.6.1) with the following options (--stranded=reverse –minaqual=10 –type=exon –-mode=intersection-nonempty) based on Gencode v33 annotations. Differential gene expression analyses were conducted using DESeq2 (v1.40.2). Extreme outliers were defined independently for each group and analyte and were removed prior to analysis. Extreme outliers were defined as values exceeding the range of three times the interquartile distance either below the first quartile (*Q1 – 3IQR*) or above the third quartile (*Q3 + 3IQR*). For differential expression comparing individuals with T21 versus D21, and B cell high/low groups, source, sex, and age were used as covariates. For visualization purposes only, *removeBatchEffect()* from the limma (v3.60.4) package was used to adjust for these same covariates.

#### Analysis of SomaScan plasma proteomics data

Data in the SomaScan adat file format were processed in R using the SomaDataIO package (v3.1.0). Extreme outliers were defined independently for each group and analyte and were removed prior to analysis. Extreme outliers were defined as values exceeding the range of three times the interquartile distance either below the first quartile (*Q1 – 3IQR*) or above the third quartile (*Q3 + 3IQR*). Differential abundance analysis was carried out in R by linear regression using the *lm()* function from the stats package (v4.4.0). For differential abundance comparing individuals with T21 versus D21, and B cell high/low groups, log_2_-transformed relative abundance served as the outcome/predictor variable and T21 status or B cell groups served as the predictor/independent variable, with age, sex and sample source as covariates. For visualization purposes only, *removeBatchEffect()* from the limma (v3.60.4) package was used to adjust for these same covariates.

#### Analysis of BCR and TCR repertoire sequencing data

Raw FASTQ files were analyzed using the Cogent NGS Immune Profiler pipeline (CogentAP), which performs quality filtering, immune receptor assembly, clonotype identification, and V(D)J annotation to generate AIRR-compliant repertoire reports and clone-level summary tables. Immune Profiler outputs included receptor sequences, CDR3 nucleotide and amino acid sequences, V(D)J gene calls, constant region assignments, and clonotype read counts for each sample.

Clone-level outputs were converted into Change-O-compatible databases and analyzed using the Immcantation framework in R (v4.4). Individual BCR isotype and TCR chain repertoires containing fewer than 30 clonotypes were excluded from downstream analyses. Repertoire diversity was quantified with the alakazam package using Hill diversity profiles, including species richness (q = 0), Shannon diversity (q = 1), and Shannon evenness with 200 bootstrap replicates. Diversity estimates were calculated using uniform sampling to account for differences in sequencing depth between repertoires. Clonal expansion (top five clone frequency) was quantified as the cumulative frequency of the five most abundant clonotypes within each repertoire. V– and J-gene usage frequencies were quantified using the Change-O package and expressed as abundance-weighted clonotype frequencies. BCR repertoires were analyzed separately for each immunoglobulin isotype (IgM, IgD, IgG, IgA, and IgE), while TCR repertoires were analyzed separately for α– and β-chain repertoires.

For somatic hypermutation (SHM) analyses, BCR repertoires were reannotated using IgBLAST against IMGT germline reference sequences to generate nucleotide-level germline alignments. SHM frequencies were calculated by comparing each rearranged V-region sequence with its inferred germline allele. Repertoire-level SHM metrics included the median mutation frequency, 90th and 95th percentile mutation frequencies, and the proportion of sequences with mutation frequencies exceeding 1%, 2%, 5%, and 10%.

#### Pathway analysis

Results from of whole blood RNA-seq and SomaScan plasma proteomics differential abundance analyses were assessed for enrichment of hallmark gene signatures using Gene Set Enrichment Analysis (GSEA)^87^. GSEA was performed in R using the fgsea package (v1.14.0) with Hallmark gene sets. Genes and proteins were ranked on log_2_-transformed fold changes (for RNA-seq) or log_2_(fold change) multiplied by −log_10_(*P*value) (for SomaScan proteomics).

#### Analysis of Xenium spatial transcriptomics

Data pre-processing, statistical analyses, and visualization were conducted in R (R 4.5+, RStudio 2025+, Bioconductor 3.21+) as detailed below. Following onboard segmentation using the Xenium Explorer 4, the data output, including cell-feature matrices, transcript coordinates, and cell boundaries were imported into the Seurat (v5.3.1) analytical framework. The dataset was filtered to exclude low-quality cells (<100 detected transcripts) and normalized using SCTransform. The spacexr package (v2.2.1) was used to perform Robust Cell Type Deconvolution to assign biological identities to the segmented cells using the HCATonsilData (v1.6.0) package single cell dataset as the reference database and using *first_type* results for cell identity and assigning niche boundaries. T-tests were used to compare cell type composition across groups as a proportion of all cells in a sample or all cells in a sample niche. Cell types were aggregated into pseudobulk datasets using Seurat and differential expression analysis was performed using the DESeq2 package (v4.5.0). Raw counts were filtered to remove low-expression genes (minimum 0.5 counts per million in half of samples). Genes were considered differentially expressed if they exhibited a Benjamini-Hochberg adjusted p-value < 0.1. Plots were generated primarily using ggplot2 (v4.0.1). Sina plots comparing cell type composition or normalized gene counts were created with the ggplot2 and the geom_sina() function from ggforce (v0.5.0). Heatmaps were generated using the tidyheatmap (v1.13.1) package. Color palettes were sourced from RColorBrewer (v1.1.3).

## DATA AVAILABILITY

All data needed to evaluate the conclusions in the paper are present in the paper, the Supplementary Tables, and the data repositories listed below. The demographics and clinical data used in this study have been deposited on the Synapse data sharing platform under accession code syn31488784 (https://doi.org/10.7303/syn31488784). The whole blood RNA-seq data used in this study have been deposited on Synapse under accession code syn31488780 (https://doi.org/10.7303/syn31488780) and Gene Expression Omnibus under accession code GSE190125 (https://www.ncbi.nlm.nih.gov/geo/query/acc.cgi?acc=GSE190125). The SomaScan plasma proteomics data used in this study have been deposited on Synapse under accession code syn31488781 (https://doi.org/10.7303/syn31488781). The BCR-seq and TCR-seq data used in this study have been deposited on the INCLUDE Data Hub under the Human Trisome Project study page (https://doi.org/10.71738/p0a9-2v09). Databases used in the generation of these results the Human Molecular Signatures Database (MSigDB, https://www.gsea-msigdb.org/gsea/msigdb/collections.jsp), the Genome Reference Consortium Human Build 38 (GRCh38, https://www.ncbi.nlm.nih.gov/grc/human), GENCODE Release 33 annotations (https://www.gencodegenes.org/human/release_33.html), and IMGT®, the international ImMunoGeneTics information system (https://www.imgt.org). Biospecimens are available through the Human Trisome Project Biobank and can be requested online through www.trisome.org.

## ACKNOWLEDGEMENTS

We thank all self-advocates with DS and their families for participation in the Human Trisome Project and all team members who contributed to this cohort study over the years. We thank Dr. E. Clambey and his team at the Flow Cytometry Shared Resource and Dr. Kim Jordan and her team at the Human Immune Monitoring Shared Resource for outstanding service in generation of the mass cytometry dataset. We are also grateful to the Colorado Translational and Sciences Institute and the Colorado Multiple Institutional Review Board for assistance in all clinical research projects involving the Crnic Institute. Special thanks to Michelle Sie Whitten and the team at the Global Down Syndrome Foundation for logistical support at multiple stages of the project. This work was supported by the NIH INCLUDE Project at the NIH Office of the Director through grants from the National Institute of Allergy and Infectious Diseases (NIAID, R01AI150305, PI: Espinosa), the National Center for Advancing Translational Sciences (NCATS, 5UL1TR002535-02S1, PI: Sokol), the National Heart, Lung and Blood Institute (NHLBI, R01HL155691-01, contact PI: M. Verneris), and the National Institute on Aging (NIA, U24AG092191, contact PI: Espinosa); NIH grant P30CA046934 (support of core facilities); the Linda Crnic Institute for Down Syndrome, the Global Down Syndrome Foundation, the Anna and John J. Sie Foundation, GI & Liver Innate Immune Program (J.M.E.), Human Immunology and Immunotherapy Initiative (J.M.E.), and the University of Colorado School of Medicine. The findings in this manuscript do not reflect the opinions of the funding agencies.

## AUTHOR CONTRIBUTIONS

M.G.D. contributed to conceptualization, analysis, methodology, investigation, and writing – original and review/editing.

N.P.E. contributed to analysis, methodology, investigation, and writing – review/editing.

B.F.N. contributed to conceptualization, analysis, methodology, investigation, and writing – original and review/editing.

E.W. contributed to conceptualization, analysis, methodology, investigation, and writing – original and review/editing.

E.H. contributed to analysis, methodology, investigation, and writing – review/editing.

N.J. contributed to conceptualization, analysis, methodology, investigation, and writing – original and review/editing.

E.C.B. contributed to methodology, investigation, and writing – review/editing.

B.B.G contributed to methodology, investigation, and writing – review/editing.

J.J.F. contributed to methodology, investigation, and writing – review/editing.

G.D. contributed to methodology, investigation, and writing – review/editing.

B.W.H. contributed to methodology, investigation, and writing – review/editing.

N.R.F. contributed to methodology, investigation, and writing – review/editing.

D.J. contributed to methodology, investigation, and writing – review/editing.

Z.A. contributed to methodology, investigation, and writing – review/editing.

A.L.R. contributed to conceptualization, methodology, investigation, funding acquisition, supervision, and writing – review/editing.

K.D.S. contributed to conceptualization, methodology, investigation, funding acquisition, supervision, and writing – review and editing.

M.D.G. contributed to conceptualization, analysis, methodology, investigation, funding acquisition, supervision, and writing – review/editing.

M.R.V. contributed to conceptualization, methodology, investigation, funding acquisition, supervision, and writing – review/editing.

J.M.E. contributed to conceptualization, methodology, investigation, funding acquisition, supervision, project administration, and writing – original and review/editing.

## COMPETING INTERESTS

J.M.E. has provided consulting services to Eli Lilly and Co., Gilead Sciences Inc., Biohaven Pharmaceuticals, Perha Pharmaceuticals, and AbbVie. All other authors declare that they have no competing interests.

## SUPPLEMENTARY TABLE LEGENDS

**Supplementary Table 1**

**a** Cell cluster annotations with key defining markers and abundances as a percentage of total B cells

**b** Reagents used for CyTOF staining.

**c** Results from linear regressions of B cell marker expression comparing individuals with trisomy 21 (T21) vs. euploid controls (D21). Results show per B cell marker the log_2_(fold-change), p-value, and adjusted p-value (q-value). Multiple hypothesis correction was performed using the Benjamini-Hochberg method. **d** Results from Differential Expression Sliding Window Analysis (DESWAN) of the effect of T21 on B cell marker expression. Differential abundance per window was determined by linear regression comparing T21 vs. D21 within each window. Columns report log_2_(fold-change) and adjusted p-value (padj) for each window. Multiple hypothesis correction was performed using the Benjamini-Hochberg method.

**e** Spearman correlation analysis between B cell marker expression in individuals with T21. Results show Spearman rho, p-value, and adjusted p-value (q-value) per B cell marker pair. Multiple hypothesis correction was performed using the Benjamini-Hochberg method.

**f** Results from beta regressions of B cell subsets comparing T21 vs. D21. Results show per subset log_2_(fold-change), p-value, and adjusted p-value (q-value). Multiple hypothesis correction was performed using the Benjamini-Hochberg method.

**g** Results from Differential Expression Sliding Window Analysis (DESWAN) of the effect of T21 on B cell subset abundance. Differential abundance per window was determined by beta regression comparing T21 vs. D21 within each window. Columns report log_2_(fold-change) and adjusted p-value (padj) for each window. Multiple hypothesis correction was performed using the Benjamini-Hochberg method.

**h** Results from beta regressions comparing phosphorylated STAT1 (pSTAT1) positivity across B cell subsets in T21 vs. D21. Results show per subset log_2_(fold-change), p-value, and adjusted p-value (q-value). Multiple hypothesis correction was performed using the Benjamini-Hochberg method.

**i** Results from Differential Expression Sliding Window Analysis (DESWAN) of the effect of T21 on pSTAT1 positivity across B cell subsets. Differential abundance per window was determined by beta regression comparing T21 vs. D21 within each window. Columns report log_2_(fold-change) and adjusted p-value (padj).

**Supplementary Table 2**

**a** Results from beta regression analyses of CyTOF immune cell profiles of live, CD45+CD66lo, B cell, and T cell populations. Comparisons include individuals with trisomy 21 (T21) with high B cell levels (B cell high) vs. euploid controls (D21); individuals with T21 with low B cell levels (B cell low) vs. D21; and T21 B cell high vs. T21 B cell low. Results show per cell type log_2_(fold-change), p-value, and adjusted p-value (q-value). Multiple hypothesis correction was performed using the Benjamini-Hochberg method.

**b** Results from DESeq2 differential expression analyses of whole-blood RNA-seq. Comparisons include T21 B cell high vs. D21; T21 B cell low vs. D21; and T21 B cell high vs. T21 B cell low. Results show per feature log_2_(fold-change), p-value, and adjusted p-value (q-value). Multiple hypothesis correction was performed using the Benjamini-Hochberg method.

**c** Results from Hallmark Gene Set Enrichment Analysis (GSEA) of whole-blood RNA-seq differential expression from (b). Results show per Hallmark gene signature the normalized enrichment score (NES), p-value, adjusted p-value (q-value), and leading edge genes. Multiple hypothesis correction was performed using the Benjamini-Hochberg method.

**d** Results from linear regression analyses of SomaScan plasma proteomics. Comparisons include T21 B cell high vs. D21; T21 B cell low vs. D21; and T21 B cell high vs. T21 B cell low. Results show per feature log_2_(fold-change), p-value, and adjusted p-value (q-value). Multiple hypothesis correction was performed using the Benjamini-Hochberg method.

**Supplementary Table 3**

**a** Results from linear regression analyses of B cell receptor (BCR) repertoire richness comparing individuals with trisomy 21 (T21) vs. euploid controls (D21) across Ig isotypes. Comparisons were made on full length and Complementarity-Determining Region 3 (CDR3) fragments. Columns report fold-change, log_2_(fold-change), p-value, and adjusted p-value (q-value). Multiple hypothesis correction was performed using the Benjamini-Hochberg method.

**b** Results from linear regression analyses of the effect of age on BCR repertoire richness. Comparisons were made on full length and Complementarity-Determining Region 3 (CDR3) fragments. Columns report fold-change, log_2_(fold-change), p-value, and adjusted p-value (q-value). Multiple hypothesis correction was performed using the Benjamini-Hochberg method.

**c** Results from linear regression analyses of BCR repertoire Shannon diversity comparing individuals with T21 vs. D21 across Ig isotypes. Comparisons were made on full length and CDR3 fragments. Columns report fold-change, log_2_(fold-change), p-value, and adjusted p-value (q-value). Multiple hypothesis correction was performed using the Benjamini-Hochberg method.

**d** Results from linear regression analyses of the effect of age on BCR repertoire Shannon diversity. Comparisons were made on full length and CDR3 fragments. Columns report fold-change, log_2_(fold-change), p-value, and adjusted p-value (q-value). Multiple hypothesis correction was performed using the Benjamini-Hochberg method.

**e** Results from linear regression analyses of BCR repertoire evenness comparing individuals with T21 vs. D21 across Ig isotypes. Comparisons were made on full length and CDR3 fragments. Columns report fold-change, log_2_(fold-change), p-value, and adjusted p-value (q-value). Multiple hypothesis correction was performed using the Benjamini-Hochberg method.

**f** Results from linear regression analyses of the effect of age on BCR repertoire evenness. Comparisons were made on full length and CDR3 fragments. Columns report fold-change, log_2_(fold-change), p-value, and adjusted p-value (q-value). Multiple hypothesis correction was performed using the Benjamini-Hochberg method.

**g** Results from beta regression analyses of BCR repertoire top five clone frequency comparing individuals with T21 vs. D21 across Ig isotypes. Comparisons were made on full length and CDR3 fragments. Columns report fold-change, log_2_(fold-change), p-value, and adjusted p-value (q-value). Multiple hypothesis correction was performed using the Benjamini-Hochberg method.

**h** Results from beta regression analyses of the effect of age on BCR repertoire top five clone frequency. Comparisons were made on full length and CDR3 fragments. Columns report fold-change, log_2_(fold-change), p-value, and adjusted p-value (q-value). Multiple hypothesis correction was performed using the Benjamini-Hochberg method.

**i** Results from linear regression analyses of BCR repertoire V gene usage comparing individuals with T21 vs. D21 across Ig isotypes. Comparisons were made on full length fragments. Columns report fold-change, log_2_(fold-change), p-value, and adjusted p-value (q-value). Multiple hypothesis correction was performed using the Benjamini-Hochberg method.

**j** Results from linear regression analyses of BCR repertoire J gene usage comparing individuals with T21 vs. D21 across Ig isotypes. Comparisons were made on full length fragments. Columns report fold-change, log_2_(fold-change), p-value, and adjusted p-value (q-value). Multiple hypothesis correction was performed using the Benjamini-Hochberg method.

**k** Results from two-sided Wilcoxon rank-sum tests (Mann–Whitney U tests) of BCR clonotype mean CDR3 length comparing individuals with T21 vs. D21 across Ig isotypes. Comparisons were made on full length fragments. Columns report Wilcoxon rank-sum test statistic (Statistic, W), effect size, adjusted p-value (q-value), and magnitude of effect size. Multiple hypothesis correction was performed using the Benjamini-Hochberg method.

**l** Results from beta regression analyses of BCR repertoire mean somatic hypermutation (SHM) frequency comparing T21 vs. D21 across Ig isotypes. Comparisons were made on full length fragments. Columns report fold-change, log_2_(fold-change), p-value, and adjusted p-value (q-value). Multiple hypothesis correction was performed using the Benjamini-Hochberg method.

**m** Results from BCR SHM tail tests comparing T21 vs. D21 across immunoglobulin (Ig) isotypes using two-sided Wilcoxon rank-sum analyses (Mann–Whitney U tests). Results are shown separately for each Ig isotype and SHM metric, including the proportion of clones with mutation frequencies exceeding 1% (prop_gt1), 2% (prop_gt2), 5% (prop_gt5), and 10% (prop_gt10), as well as the 90th (q90) and 95th (q95) percentile SHM frequencies within each repertoire. Columns report the compared groups, sample sizes (n1, n2), Wilcoxon test statistic, nominal P value, and Benjamini–Hochberg-adjusted q value (BHadj_pval).

**Supplementary Table 4**

**a** Results from linear regression analyses of T cell receptor (TCR) repertoire richness comparing individuals with trisomy 21 (T21) vs. euploid controls (D21) across TCR alpha and beta chains. Comparisons were made on Complementarity-Determining Region 3 (CDR3) fragments. Columns report fold-change, log_2_(fold-change), p-value, and adjusted p-value (q-value). Multiple hypothesis correction was performed using the Benjamini-Hochberg method.

**b** Results from linear regression analyses of the effect of age on TCR repertoire richness. Comparisons CDR3 fragments. Columns report fold-change, log_2_(fold-change), p-value, and adjusted p-value (q-value). Multiple hypothesis correction was performed using the Benjamini-Hochberg method.

**c** Results from linear regression analyses of TCR repertoire Shannon diversity comparing individuals with T21 vs. D21 across TCR alpha and beta chains. Comparisons CDR3 fragments. Columns report fold-change, log_2_(fold-change), p-value, and adjusted p-value (q-value). Multiple hypothesis correction was performed using the Benjamini-Hochberg method.

**d** Results from linear regression analyses of the effect of age on TCR repertoire Shannon diversity. Comparisons were CDR3 fragments. Columns report fold-change, log_2_(fold-change), p-value, and adjusted p-value (q-value). Multiple hypothesis correction was performed using the Benjamini-Hochberg method.

**e** Results from linear regression analyses of TCR repertoire evenness comparing individuals with T21 vs. D21 across TCR alpha and beta chains. Comparisons were made on CDR3 fragments. Columns report fold-change, log_2_(fold-change), p-value, and adjusted p-value (q-value). Multiple hypothesis correction was performed using the Benjamini-Hochberg method.

**f** Results from linear regression analyses of the effect of age on TCR repertoire evenness. Comparisons were made on CDR3 fragments. Columns report fold-change, log_2_(fold-change), p-value, and adjusted p-value (q-value). Multiple hypothesis correction was performed using the Benjamini-Hochberg method.

**g** Results from beta regression analyses of TCR repertoire top five clone frequency comparing individuals with T21 vs. D21 across TCR alpha and beta chains. Comparisons were made on CDR3 fragments. Columns report fold-change, log_2_(fold-change), p-value, and adjusted p-value (q-value). Multiple hypothesis correction was performed using the Benjamini-Hochberg method.

**h** Results from beta regression analyses of the effect of age on TCR repertoire top five clone frequency. Comparisons were made on CDR3 fragments. Columns report fold-change, log_2_(fold-change), p-value, and adjusted p-value (q-value). Multiple hypothesis correction was performed using the Benjamini-Hochberg method.

**i** Results from linear regression analyses of TCR repertoire V gene usage comparing individuals with T21 vs. D21 across TCR alpha and beta chains. Comparisons were made on full length fragments. Columns report fold-change, log_2_(fold-change), p-value, and adjusted p-value (q-value). Multiple hypothesis correction was performed using the Benjamini-Hochberg method.

**j** Results from linear regression analyses of TCR repertoire J gene usage comparing individuals with T21 vs. D21 across TCR alpha and beta chains. Comparisons were made on full length fragments. Columns report fold-change, log_2_(fold-change), p-value, and adjusted p-value (q-value). Multiple hypothesis correction was performed using the Benjamini-Hochberg method.

**k** Spearman correlation analysis between BCR and TCR repertoire metrics in individuals with T21. Results show Spearman rho, p-value, and adjusted p-value (q-value) per metric pair. Multiple hypothesis correction was performed using the Benjamini-Hochberg method.

**Supplementary Table 5**

**a** Results from Student’s t-tests comparing in individuals with trisomy 21 (T21) vs. euploid controls cell composition at the tonsil level. Columns report per cell type sample sizes, means, standard deviations, mean difference, confidence intervals, fold-change, log_2_(fold-change), p-value, and t-statistic.

**b** Results from Student’s t-tests comparing in individuals with trisomy 21 (T21) vs. euploid controls cell composition at the germinal center (GC) niche level. Columns report per cell type sample sizes, means, standard deviations, mean difference, confidence intervals, fold-change, log_2_(fold-change), p-value, and t-statistic.

**c** Results from Student’s t-tests comparing in individuals with trisomy 21 (T21) vs. euploid controls cell composition at the mantle zone (MZ) niche level. Columns report per cell type sample sizes, means, standard deviations, mean difference, confidence intervals, fold-change, log_2_(fold-change), p-value, and t-statistic.

**d** Results from Student’s t-tests comparing in individuals with trisomy 21 (T21) vs. euploid controls cell composition at the epithelium (Epi) niche level. Columns report per cell type sample sizes, means, standard deviations, mean difference, confidence intervals, fold-change, log_2_(fold-change), p-value, and t-statistic.

**e** Results from Student’s t-tests comparing in individuals with trisomy 21 (T21) vs. euploid controls cell composition at the T cell zone (TCZ) level. Columns report per cell type sample sizes, means, standard deviations, mean difference, confidence intervals, fold-change, log_2_(fold-change), p-value, and t-statistic.

**Supplementary Table 6**

**a** Results from DESeq2 of tonsil spatial transcriptomics comparing individuals with trisomy 21 (T21) vs. euploid controls (D21). Columns display per cell type (Clusterid) and gene (Gene_name, Geneid) the fold-change, log_2_(fold-change), p-value, and adjusted p-value (padj). Multiple hypothesis correction was performed using the Benjamini-Hochberg method.

## Notes

https://doi.org/10.7303/syn31488784

https://doi.org/10.7303/syn31488780

https://www.ncbi.nlm.nih.gov/geo/query/acc.cgi?acc=GSE190125

https://doi.org/10.7303/syn31488781

https://doi.org/10.71738/p0a9-2v09

